# Microglia-triggered hyperexcitability in the cerebellum depresses animal behaviors

**DOI:** 10.1101/353730

**Authors:** Masamichi Yamamoto, Minsoo Kim, Hirohiko Imai, Gen Ohtsuki

**Author notes:** These authors contributed equally (Masamichi Yamamoto, Minsoo Kim). Correspondence: Gen Ohtsuki, Ph.D.

## Abstract

Clinical studies have suggested that cerebellar dysfunction is involved in various psychiatric disorders, including autism spectrum disorders, dyslexia, and depressive disorders. However, the physiological aspect is less-advanced. Here, we comprehensively investigated the immune-triggered excitability plasticity in the cerebellum. Activated microglia (MG) via exposure to bacterial endotoxin lipopolysaccharide or heat-killed Gram-negative bacteria induced a potentiation of the excitability of Purkinje neurons, which was suppressed by MG-activity inhibitor and MG-depletion. An inflammatory cytokine, tumor necrosis factor-α (TNF-α) released from MG triggered this plasticity. While our new two-photon FRET ATP-imaging showed an increase in ATP concentration following endotoxin exposure, both TNF-α and ATP secretion facilitated synaptic transmission. Inflammation in the cerebellar anterior vermis *in vivo* immobilized animals, and reduced sociability. Such abulia-like behavioral impairments were reverted by TNF-α-inhibition and MG-depletion. Resting-state functional MRI revealed overconnectivity between the inflamed cerebellum and prefrontal neocortical regions, which may underlie the psychomotor depressiveness in animals.

## Main text

Excessive activation of microglia (MG)—the resident immune cells of the central nervous system (CNS)—causes neuroinflammatory responses following immune challenges in the brain. While accumulating evidence prevailed a notion that MG are associated with not only inflammation consisting of “tumor, rubor, calor et dolor” due to microbial pathogens, but also emotional or mood related psychiatric diseases accompanied by psychological stress or morbidity, the full physiological effects of immune-related responses on the CNS remain unclear^1–9^.

Recent reports showed that transient exposure of hippocampal slices to the Gram-negative bacterial endotoxin, lipopolysaccharide (LPS), enhances presynaptic excitatory transmitter release^10^ and depresses the postsynaptic efficacy of glutamate receptors under conditions of low oxygenation^11^, through MG activation. While activated MG release ATP and reactive oxygen species resulting in the induction of these synaptic plasticity^10,11^, inflammatory cytokines are also known to modulate synaptic transmission^12–15^. In contrast, CNS neurons exhibit a form of plasticity that enables them to modify intrinsic firing properties in response to the external activity. Several reports showed that activated MG may alter non-synaptic membrane excitability in neurons in the retina, hippocampus, or neocortex; however, direct evidence regarding the long-term plasticity of intrinsic excitability via MG, the mechanisms at cellular level, and the relevance to animal diseases are yet lacking.

In the present study, we investigated whether MG, activated by exposure to bacterial endotoxin or heat-killed bacteria, modulate cerebellar neuronal excitability, and we aimed to elucidate the induction mechanism of excitability plasticity. Further, we examined the effect of aberrant neuronal excitability on animal behavior and functional connectivity during cerebellar inflammation, and we challenged whether the symptoms of acute cerebellitis could be rescued.

## Results

### Microglia modulate the intrinsic excitability of cerebellar Purkinje neurons

To investigate the nature of immune-neuronal interactions, we examined the neuronal excitability of Purkinje cells—the principle output neurons of the cerebellar cortex—following exposure to the endotoxin LPS (10-12 μg/mL), an outer membrane component of Gram-negative bacteria. LPS was applied to cerebellar slices in a bath-chamber, after which the firing properties of Purkinje neurons were examined under current-clamp (**Fig. 1a,b**). The firing frequency of neurons in response to depolarization with different current pulses was significantly higher after LPS application than that under control conditions. To investigate the time course of firing changes, we subjected slices to transient LPS exposure, then continuously monitored subsequent firing properties. Firing frequency was significantly higher relative to baseline following LPS exposure than control (*p < 0.001) (**Fig. 1c,d**). We next applied heat-killed Gram-negative bacteria (**Fig. 1e**). Neurons treated with heat-killed *Escherichia coli* (*E. coli*) *0111:B4* (HKEB) or *Pseudomonas aeruginosa* (HKPA) (10^7^ cells/mL) exhibited significantly higher firing frequencies than control neurons. However, neurons exposed to Gram-positive bacteria (heat-killed *Streptococcus pneumoniae*, HKSP) that have no LPS did not exhibit any significant increases in firing. Results obtained from long-term recordings also indicated the firing increase following exposure to heat-killed Gram-negative bacteria (**Fig. 1f,g**) (HKEB and HKPA, *p < 0.03) but not Gram-positive bacteria (HKSP, p > 0.1). Action potential waveforms also suggested that Purkinje neurons showed enhanced excitability following endotoxin exposure (**Supplementary Table 1**). Another major cell-wall component derived from *E coli,* peptidoglycan (PGN), did not significantly alter the waveforms (Supplementary Table 1). Experiments with NBQX suggested that this firing plasticity was induced without the involvement of AMPA receptors (*p < 0.05) (**Supplementary Fig. 1a,b**). Thus, our findings suggest that endotoxin exposure substantially altered the intrinsic membrane excitability of neurons.

**Figure 1.**
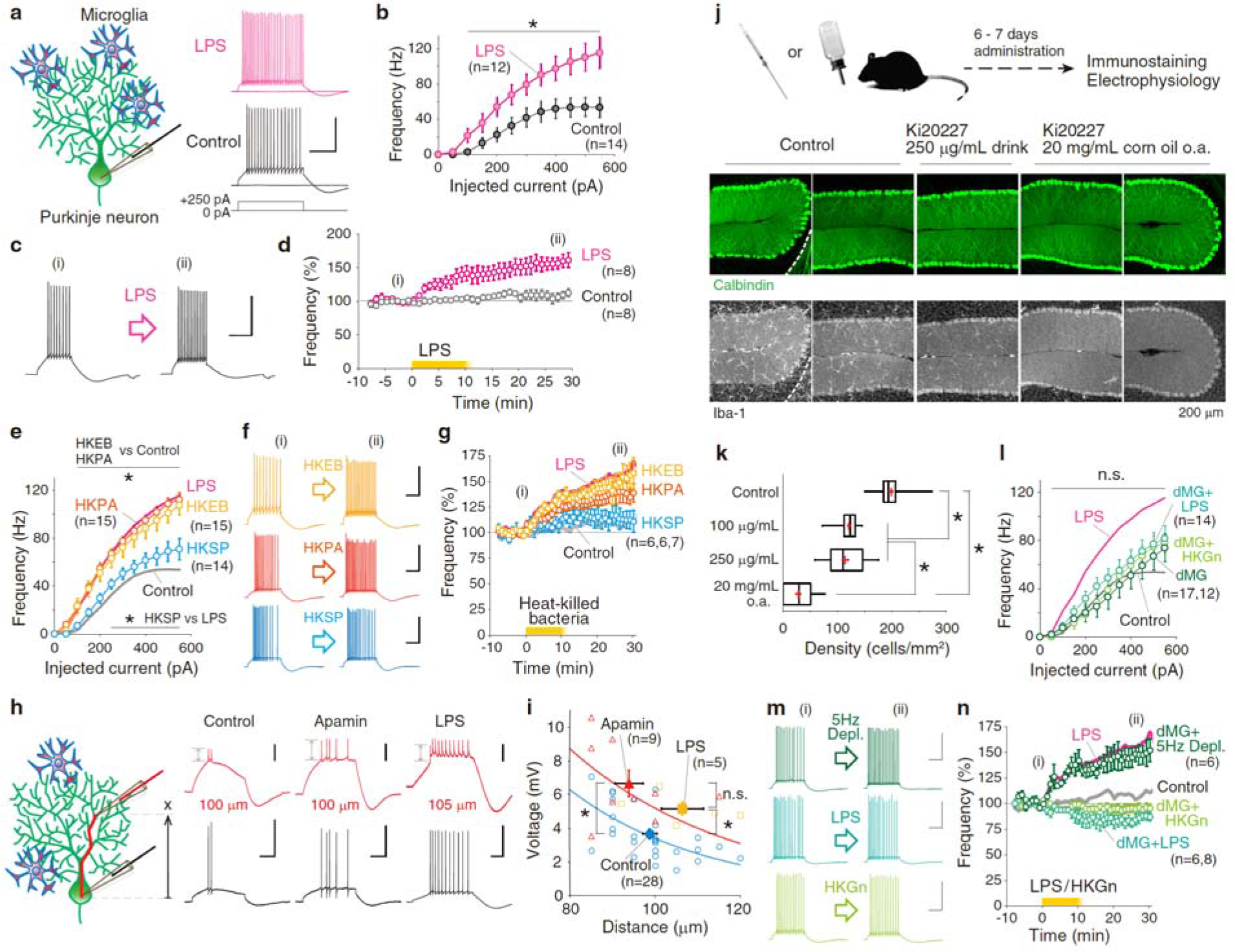
Microglia activation alters excitability in cerebellar neurons. **a,** Representative action potential (AP) firings of control and LPS (10-12 μg/mL)-treated Purkinje neurons in response to depolarization pulses. **b,** Increase in the firing frequency following bath-application of LPS. **c, d,** Representative AP firing before and after LPS exposure ((i) and (ii), respectively), and time courses of the normalized frequency. Representative AP traces in (c) were obtained from the corresponding time points in (d). **e,** Bath-application of heat-killed Gram-negative (10^7^ cells/ml, HKEB and HKPA, respectively) but not Gram-positive (HKSP) bacteria increases firing frequency. **f, g,** Representative firing and normalized time courses of long-lasting recordings of HKEB, HKPA and HKSP application. **h,** Representative AP firings from soma (black) and dendrite (red). Distance of dendrite patching (X) is shown below each trace. **i,** Distance-voltage plot of back-propagated APs. **j,** Depletion of microglia (MG) by Ki20227-administration for a week. o.a.: oral administration. **k,** Density of MG of control and Ki20227-administered cerebella (*p < 0.05, Kruskal-Wallis test). **l,** Suppression of the increase in firing frequency by exposure to LPS and HKGn (heat-killed Gram negative bacteria mixture: HKEB+HKPA). **m, n,** Impairment of the excitability increase by exposure to LPS and HKGn in the MG-depleted cerebella. Time courses of the change in firing frequency normalized between −5 to −1 min in (d), (g) and (n). Reagents-application began at 0 min and continued for 10 min. Scale: 40 mV and 200 ms (a, c, f and m), and 5 mV, 40 mV and 100 ms (h).

Previously, Belmeguenai *et al.*^16^ demonstrated that Purkinje cells exhibit intrinsic plasticity (IP), which is defined as an increase in firing frequency, with shortening spike pauses, lasting for more than 30 min^16–19^. Simultaneous somato-dendritic recordings demonstrated the enhancement of dendritic excitability accompanied with IP^17^. These increases in firing frequency and dendritic excitability are induced by transient depolarization of the neuronal membrane or parallel fiber stimulation via phosphatase-dependent signaling and down-regulation of small conductance Ca^2+^ activated K^+^ channels (SK channels). Findings from conditional knockout mice for protein phosphatase 2B (PP2B) suggest that the IP is associated with cerebellar motor coordination^20^. Considering results obtained from occlusion experiments of the IP by pre-exposed LPS (**Supplementary Fig. 1c**) and impairment of IP induction under SK channel blockade by apamin (**Supplementary Fig. 1d**), the firing increase following endotoxin exposure was induced through the same molecular signaling for IP^16–19^, depending on intra-neuronal Ca^2+^ and protein phosphatases PP1, PP2A, and PP2B (**Supplementary Fig. 1e-g**).

We next investigated changes in action potentials after LPS-application on both soma and dendrites (**Fig. 1h,i**). We applied somatic depolarization pulses under control condition, in the presence of an SK-channel blocker, and after LPS-exposure. Passively back-propagated action potentials toward dendrites significantly increased in amplitude following LPS-administration (*p < 0.04), relative to those under SK channel blockade (*p < 0.002), in a distance-dependent manner along soma-dendrite axis^17^ (**Fig. 1i**), suggesting that MG activation by endotoxin enhances excitability at dendrites via down-regulation of SK2 channels.

Given that the increase in firing frequency was triggered by MG, elimination of MG may block this plasticity. To deplete MG in living animals, we administered the colony-stimulating factor 1 receptor (CSF-1R) kinase inhibitor Ki20227^21,22^. Following continuous oral administration of Ki20227 or with drink in three different concentration, the number of MG in the immunostained cerebellar cortex was reduced to 14% at maximum (*p < 0.00001, *multiple comparison*) (**Fig. 1j,k** and **Supplementary Fig. 2**). Therefore, our Ki20227-administration depleted almost all the MG in the cerebellum. When the neuronal excitability was examined in the MG-depleted cerebella, no increases in firing was observed against exposure to LPS and heat-killed Gram-negative bacteria (HKGn: HKEB+HKPA) (**Fig. 1l**). Long-term recordings also showed that, while innate firing frequency increased after 5-Hz conditioning for IP (*p < 0.05), exposure to LPS and HKGn no longer increased the frequency (LPS, *p < 0.05 of reduction; HKGn, p > 0.3) (**Fig. 1m,n**). These results indicate that MG are involved in increasing the excitability of cerebellar neurons, which is supported by results of minocycline co-application (p > 0.7) (**Supplementary Fig. 3**).

### Involvement of inflammatory cytokine in the excitability plasticity

A possible mechanism for MG-triggered neural hyperexcitability is mediated by inflammatory cytokines, including TNF-α, interleukin (IL)-6, and IL-1β. Among them, TNF-α is released in the earliest phase of immune-cell stimulation. We first examined the firing frequency of neurons treated with TNF-α. Firing frequency of TNF-α-treated neurons was significantly higher than in control or PGN-treated (**Fig. 2a,b**). In addition, co-application of LPS or HKGn bacteria with the TNF-α inhibitor C87 abolished the increase in firing frequency (both p > 0.5) (**Fig. 2c**). To confirm the effect of TNF-α, neurons were subjected to bath-application of TNF-α, and the TNF-α treatment increased firing frequency similar to that observed following LPS administration (*p < 0.03) (**Fig. 2d**). Application of LPS to Purkinje neurons pre-exposed to TNF-α prevented the excitability increase (p > 0.5), implying that TNF-α release and excitability plasticity were both resulted from LPS application. Although our results suggest that TNF-α directs Purkinje neurons to enhance excitability, the mechanisms underlying this process remain unclear. A series of studies in hippocampal neurons previously demonstrated that TNF-α activates phosphatase signaling via TNF receptor 1 in CA1 pyramidal neurons^12,13^. Then, we subjected Purkinje neurons to bath-application of TNF-α under suppression of phosphatase activity by intra-neuronal okadaic acid, but no increases were observed in firing frequency (p > 0.3) (**Fig. 2e**), suggesting the TNF-α-mediated firing plasticity occurred through phosphatase activity.

**Figure 2.**
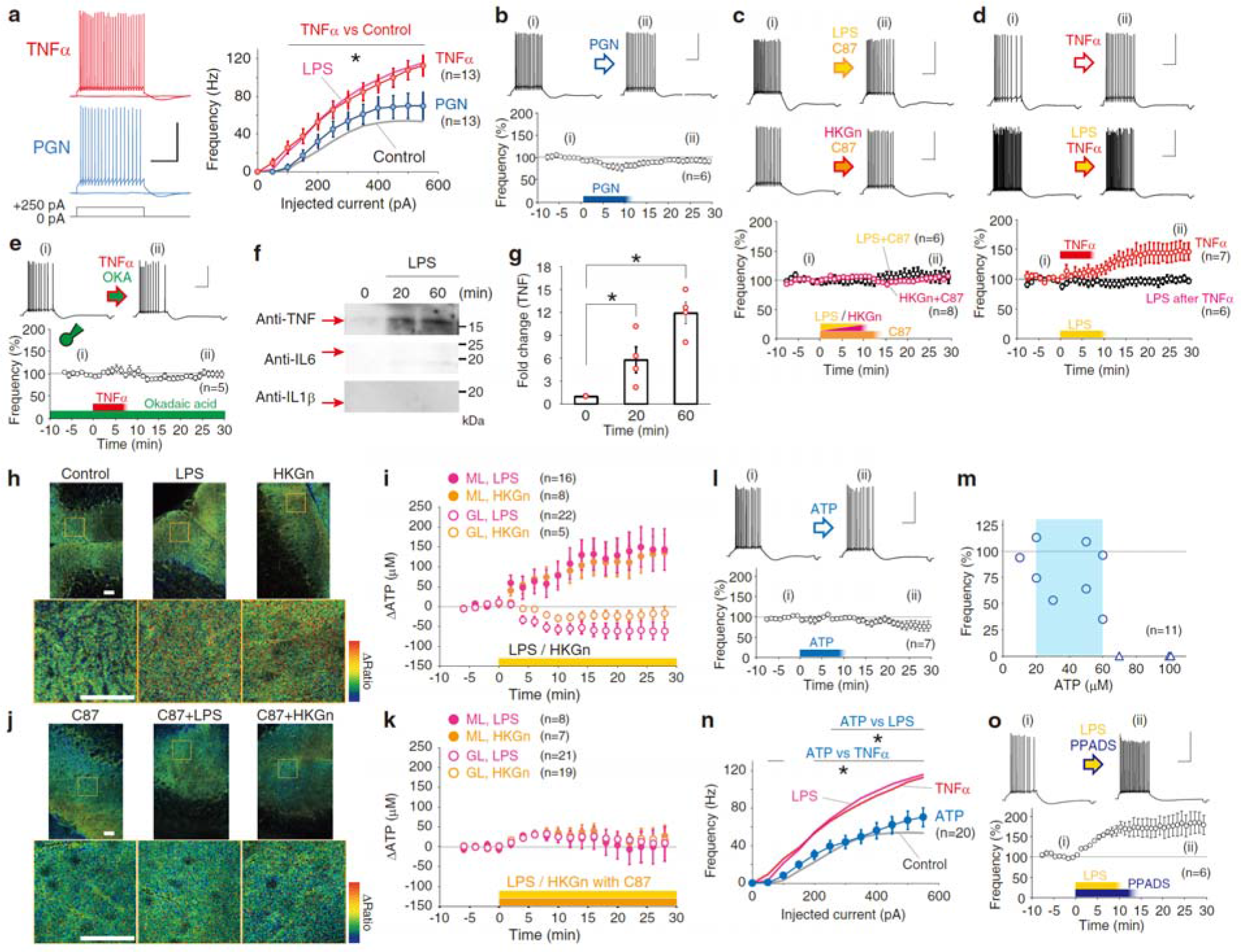
Involvement of microglial mediators in the modulation of neuronal excitability. **a,** Firing frequency of Purkinje neurons exposed to TNF-α (100 ng/mL) and PGN (10 μg/mL) against depolarization pulses (*p < 0.03). Mean firing frequency of LPS-exposed and control neurons are merged for comparison. **b,** PGN-exposure. **c,** Suppression of LPS- and HKGn-induced firing plasticity by TNF-α inhibitor, C87 (40 μM). **d,** TNF-α bath-application increased the firing frequency and occluded the effect of LPS. **e,** TNF-α-induced firing frequency increase was abolished by the intra-neuronal okadaic acid (150 nM). **f,** Western blotting of TNF, IL-6 and IL-1β protein in supernatant following LPS exposure for 0, 20, and 60 minutes. **g,** Fold change of TNF-level normalized by at 0 min. **h,** FRET imaging with ATP-probe, ATeam-GO2, shows the increase in ATP in the cerebellar slices. **i,** Time courses of changes in ATP concentration in the molecular layer (ML) and granule cell layer (GL). **j, k,** FRET ATP-imaging under TNF-α blocker C87. The ATP increase in ML by endotoxin was abolished by TNF-α inhibition. **l,** 20-60 μM ATP exposure. **m,** ATP-firing change plot indicates a cease of firing in high ATP concentration. **n,** Firing frequency after ATP administration. **o,** Firing increase was not abolished by P2R inhibitor, PPADS (50 μM). All right angle scales: 40 mV and 200 ms. Bar scales (h and j): 50 μm.

Western blotting confirmed the increase in the amount of TNF-α, but not IL-6 or IL-1β (**Fig. 2f,g**) by a transient exposure to LPS, indicating that TNF-α release mediates signaling between MG and Purkinje neurons. We also monitored action potential firing in the presence of protein-synthesis inhibitors anisomycin and cycloheximide, which did not prevent LPS-induced firing increases (*p < 0.03 for both) (**Supplementary Fig. 4**). These findings suggest that endotoxin exposure may stimulate TNF-α release from MG in a protein-synthesis-independent manner, and that diffusible TNF-α triggered the increase in neuronal excitability.

It is also possible that bacterial endotoxin triggers excitability plasticity via ATP release from tissues including MG^2,3,10,23^. Hence, we confirmed ATP synthesis and release in the whole cerebellum in knock-in mice with the ATeam probe GO-ATeam2 (see Methods), which we developed newly. Two-photon fluorescence/Förster resonance energy transfer (FRET) imaging suggested that continuous exposure to both LPS and HKGn increase ATP concentration prominently in the cerebellar molecular layer (**Fig. 2h,i** and **Supplementary Fig. 5a**). We estimated the ATP concentration changes from FRET ratio changes against endotoxin exposure (see Methods), and the extent of ATP increase reached 140 μM in the molecular layer (**Fig. 2i**). Increase in the ATP was prevented under TNF-α inhibition (**Fig. 2j,k**), suggesting the ATP synthesis follows TNF-α secretion. Next, we tested whether the excitability of Purkinje neurons is modulated by bath application of ATP with various concentrations (10-100 μM) (**Fig. 2l-n**). The firing frequency did not increase upon exposure to ATP (20-60 μM, p > 0.1) (**Fig. 2l,n**). Additionally, even under the purinergic receptor (P2R) inhibitor PPADS, firing frequency was increased upon exposure to LPS (*p < 0.05) (**Fig. 2o**).

### Facilitation of synaptic transmission following microglial activation

Regarding the effect of endotoxin exposure in synapses, we examined whether LPS administration alters spontaneous excitatory postsynaptic currents (sEPSCs). Results indicated no change in postsynaptic responsiveness in Purkinje neurons, except for a reduction in PPADS+LPS group (**Fig. 3a-d**). However, administration of heat-killed Gram-negative bacteria, TNF-α, or ATP produced a significant increase in sEPSC frequency (**Fig. 3e,f**). Our findings suggest that activated MG prompted vesicular release from presynaptic neurons, depending on both TNF-α and ATP. Extracellular ATP may increase in vesicular release via purinergic receptors on presynaptic neurons. In fact, in MG-depleted cerebella, there were no significant differences among MG-depleted control, LPS, and HKGn exposure groups in amplitude or in frequency, indicating the involvement of certain mediators from MG (**Supplementary Fig. 6**).

**Figure 3.**
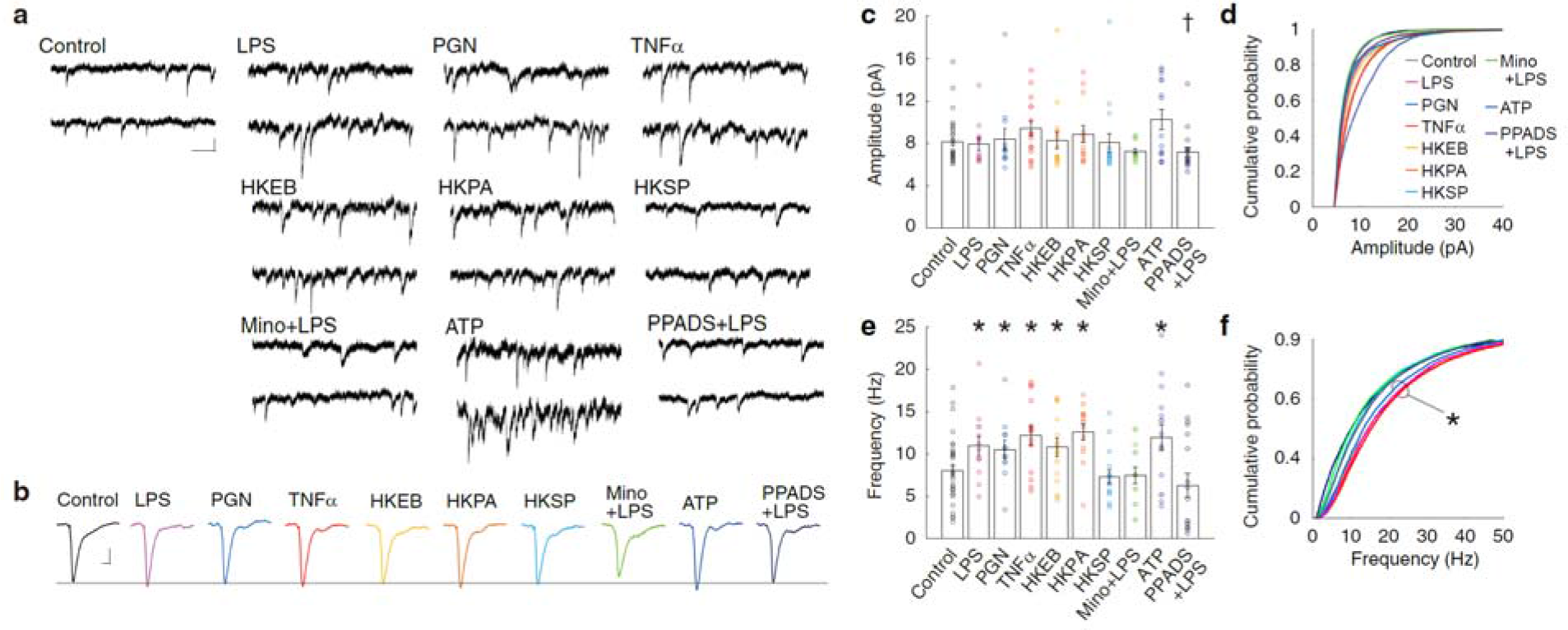
Increase in the release of excitatory synaptic transmissions following microglia activation in both TNF-α and ATP-dependent manners. **a,** Spontaneous EPSC (sEPSC) in control, LPS, PGN, TNF-α, HKEB, HKPA, HKSP, Minocycline+LPS, ATP and PPADS+LPS. Scale: 10 pA, 100 ms. **b,** Averaged representative sEPSC. Scale: 2 pA, 10 ms. **c-f,** Bar graphs (mean ± SEM) and cumulative probability graphs of amplitude (c, d) and frequency (e, f) of sEPSC in different experiments. A total of 32, 13, 12, 15, 16, 13, 16, 14, 15, and 18 cells were observed, respectively. While asterisks indicate a significance increase (Mann-Whitney *U*-test) against the control condition, a dagger indicates the decrease. Suppression of the increase in sEPSC-frequency by PPADS suggests the involvement of P2R channels.

### Endotoxin-TLR4-TNF-α signal involvement in the hyperexcitability

Next, we investigated whether the endotoxin receptors involved in MG-associated signaling. LPS is classically known to bind complement receptor 3 in immune cells^23,24^. Zhang et al.^11^ revealed that application of LPS under hypoxic conditions resulted in the release of superoxide anion via complement receptor and NADPH oxidase activity. Here, we applied LPS in the presence of apocynin, an NADPH oxidase inhibitor. However, we observed no suppression of firing increases (*p < 0.03) (**Fig. 4a**), implying that neither superoxide nor the complement receptor pathway are involved in the induction of excitability plasticity.

**Figure 4.**
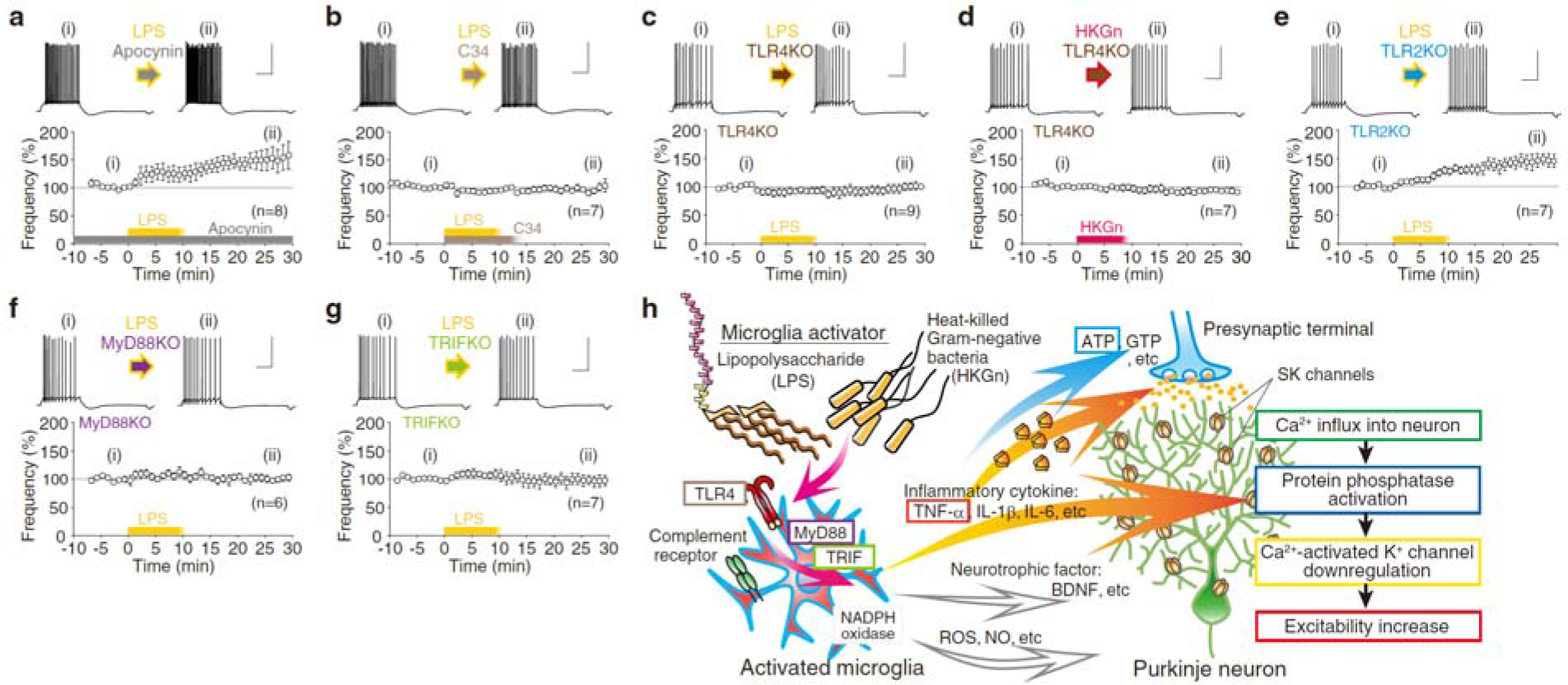
Endotoxin-TLR4-TNF-α signal involvement of the excitability increase plasticity by microglia activation. **a,** Firing increase by LPS under a nicotinamide adenine dinucleotide phosphate (NADPH)-oxidase inhibitor, apocynin (100 μM). **b,** LPS-induced firing increase plasticity was abolished by Toll-like receptor 4 (TLR4) blocker, C34 (40 μM). **c-e,** Impairment of the excitability plasticity in TLR4 KO mice but not in TLR2 KO. **f, g,** Impairment of the excitability plasticity in transgenic mice lacking TLR4 downstream proteins MyD88 and TRIF. Scales are 40 mV and 200 ms. **h,** Summary of signaling cascade.

The Toll-like receptor (TLR) family comprises pattern recognition receptors that are abundantly expressed on the surface of immune cells. The extracellular domain of leucine-rich repeats of TLR4 specifically recognize LPS^25^, and its dimerized complex is thought to trigger the intracellular signaling cascade. We examined MG signaling using pharmacology and transgenic animals. Under the suppression of TLR4 by C34, LPS exposure did not cause the change in firing frequency (p > 0.1) (**Fig. 4b**). While exposure of TLR4-knockout cerebellar slices to LPS and HKGn abolished the induction of firing increase (**Fig. 4c,d**), Purkinje neurons in TLR2-knockout mice showed an increase in firing sustainably (**Fig. 4e**), suggesting that TLR4 but not TLR2 is essential for the observed responses (**Supplementary Fig. 7a,b**). We next aimed to determine the downstream of TLR4 using knockout mice for myeloid differentiation primary response gene 88 (MyD88) and Toll/IL-1 receptor (TIR)-domain-containing adapter-inducing interferon-β (TRIF). While MyD88 is a TIR domain-containing adapter common in TLR signaling pathways, TRIF mediates the MyD88- independent pathway as the downstream of TLR3 and TLR4^24^. Both are involved in inflammatory cytokine release in macrophages and MG. In MyD88- and TRIF-knockout mice, no significant difference was observed in firing frequency between LPS-treated and untreated neurons (**Supplementary Fig. 7c,d**) nor in firing properties (**Supplementary Table 2**). In addition, LPS exposure did not produce long-lasting firing increases (both, p > 0.7) (**Fig. 4f,g**), suggesting that both pathways in MG for the excitability plasticity induction. While, in MG, the precise machinery underlying the exocytosis of inflammatory cytokines remains obscure, a certain molecule downstream of both signals may prime the secretion of soluble TNF-α, as illustrated in a summary cascade of the endotoxin-induced excitability plasticity in the cerebellum (**Fig. 4h**, **Supplementary Fig. 8** and **Supplementary Fig. 9**).

### Depressive behaviors of animals with cerebellar inflammation

To reveal the *in vivo* physiological significance of excess immune activity in the cerebellum, we observed animals’ behavior after bacterial endotoxin infusion in anterior lobes of the cerebellar vermis. Unexpectedly, we found that the spatial exploratory behavior of freely moving rats in an open field was significantly reduced by LPS (1 mg/mL)- and HKGn (HKEB+HKPA, 10^9^ cells/mL for each)-administration (**Fig. 5a,b** and **Supplementary Video 1**). In social interaction test, LPS- and HKGn-injected rats exhibited considerably less interest in siblings (**Fig. 5c**). Immobility time during a forced swim test became significantly elongated than that in control PBS-injected animals (**Fig. 5d**). These results suggest that the behavior of animals with cerebellar injection is depression like or abulic, in contrast to autistic behaviors in animals with less excitable Purkinje neurons^26,27^. To examine the extent of repetitive movements, we conducted marble burying test and found that rats with cerebellar bacterial endotoxin infusion showed less burying behavior (**Fig. 5e** and **Supplementary Fig. 10**). Animals with cerebellar injection did not show significant motor discoordination or ataxia (**Supplementary Fig. 11a,b**). And, the behavioral modulation lasted only until the following day (**Supplementary Fig. 11c**). Magnetic resonance (MR) fluid-attenuated inversion recovery (FLAIR) images clearly showed inflammation at the restricted position of the HKGn-injection (**Fig. 5f**). In addition, animals subjected to LPS in cerebellar hemispheres showed no reduction of exploratory behavior (**Supplementary Fig. 11d**). We further recorded Purkinje neuron activity following the drug injection, and we observed a significant increase in the firing frequency of neurons from LPS- and HKGn-injected cerebella (**Supplementary Fig. 12**). Taken together, hyperexcitability in the cerebellum through region-specific inflammation caused a reduction of the psychomotor activity.

**Figure 5.**
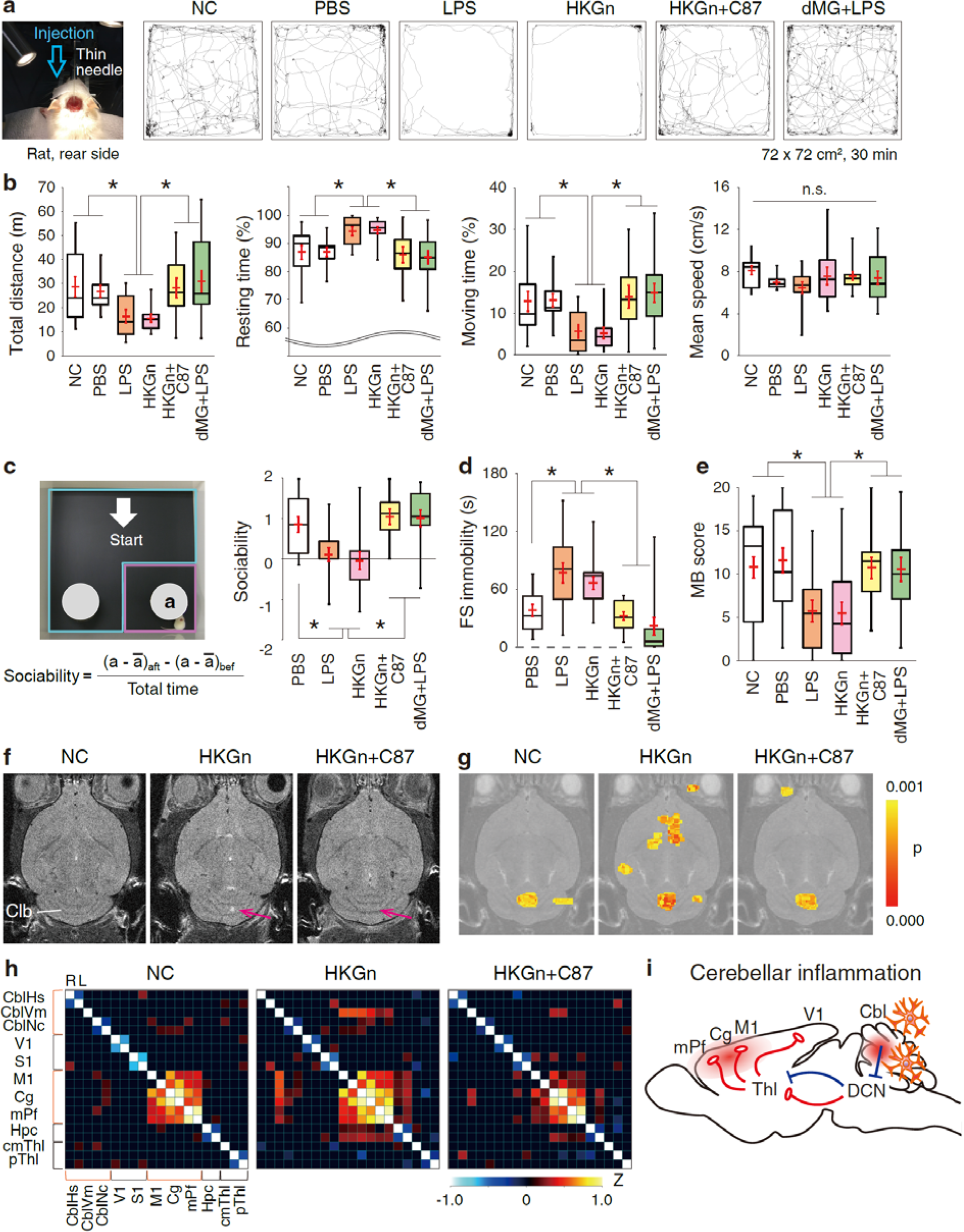
Induction and rescue of psychomotor depressiveness attributed from cerebello frontal functional overconnectivity. **a,** Representative trajectories of exploration behavior of drug infused rats in the open field arena. PBS, LPS, heat-killed Gram-negative bacteria mixture (HKGn), HKGn+C87, LPS to microglia-depleted animals (dMG+LPS) or nothing (NC) was injected into the anterior cerebellar vermis. **b,** Box plots of the total distance, resting time, moving time, and mean speed. Overlapping red marks represent mean ± SEM. Co-injection of TNF-α inhibitor C87 and MG-deletion significantly reduced the sluggishness of animals. **c,** Social interaction. **d,** Forced swim test. e, Marble burying score. *p < 0.05, *multiple comparison*. **f,** FLAIR magnetic resonance images indicate inflammation in HKGn, but not in HKGn+C87. Magenta arrows indicate the locations of injection. **g,** Resting-state functional magnetic resonance imaging (rs-*f*MRI) of control, HKGn- and HKGn+C87-infused rats. A seed was given in the anterior lobe of cerebellar vermis. Color-code indicates the functional connectivity obtained from BOLD signal. **h,** Seed-seed correlation maps shown with Z-score. Seeds were applied in the right and left hemispheres as marked by R and L, respectively, except for the anterior lobe of the cerebellar vermis (CblVm) and centro-medial thalamus (cmThl). Please note that the correlation is enhanced in HKGn between CblVm and M1+Cg+mPf (*i.e.*, primary motor cortex + cingulate cortex + medial prefrontal cortex). Abbreviations of brain regions are described in Methods. **i,** Schematic drawing of activated regions during the cerebellar inflammation.

### Recovery of behavioral disturbances by immune suppression

Provided that activated MG secrete molecules during inflammation *in vivo*, microinfusion of such molecules may suffice to modulate animal behavior. We next applied TNF-α (20 μg/mL) and ATP (20 mM) into the cerebellar anterior lobes, and found that TNF-α, but not ATP, reduced behaviors (**Supplementary Fig. 13**). This result indicates that suppression of TNF-α should help to reduce the animals’ psychomotor depressiveness. In fact, co-injections of HKGn with C87 (2-4 mM) into the anterior lobes of the cerebellar vermis showed a clear recovery of impairment of psychomotor behaviors (**Fig. 5a-e** and **Supplementary Video 1**, HKGn+C87), as well as inflammation (**Fig. 5f**, HKGn+C87) and neuronal excitability (**Supplementary Fig. 12**, HKGn+C87). Further, we speculated that the MG-depletion may also cancel the psychomotor depressiveness by endotoxin, and the results obtained in Ki20227-administered animals proved that this was really the case (**Fig. 5a-e**, dMG+LPS). Together, these striking effects of endotoxin infusion were substantially rescued by both co-injection with TNF-α inhibitor and MG-depletion (**Supplementary Fig. 14**).

In addition, to investigate the causality of the abnormal behaviors, we conducted functional MR imaging (*f*MRI) of resting-state animals, which showed a distinct enhancement of correlated signals among the HKGn-infused cerebellar anterior vermis and frontal neocortical areas, including the medial prefrontal cortex (mPf), cingulate cortex (Cg), and primary motor cortex (M1) (**Fig. 5g-i** and **Supplementary Fig. 15**). And, these functional overconnectivity caused by cerebellar inflammation (*i.e.*, hyperexcitation in the cerebellar cortex) was reverted by the TNF-α inhibition (**Fig. 5g,h**, HKGn+C87), suggesting that the modulation of the long-range connectivity between the cerebellar vermis and prefrontal areas may relate to the animal behavioral impairments.

## Discussion

The results of the present study indicate that activated MG elicit the induction of long-term potentiation of intrinsic excitability in cerebellar Purkinje neurons. We observed that both pharmacological suppression and depletion of MG activity by a CSF1 receptor inhibitor (Ki20227) and minocycline abolished firing increases by exposure to endotoxin or heat-killed Gram-negative bacteria, suggesting that MG are involved in the induction of the excitability plasticity (**Fig. 1m,n** and **Supplementary Fig. 3**). And, our results suggest that endotoxin-induced plasticity of the intrinsic excitability share molecular signaling via down-regulation of SK channels in neurons (**Supplementary Fig. 1** and **Fig. 1h,i**). To our knowledge, the present study is the first to demonstrate increases in excitability in soma and dendrites following transient exposure of CNS neurons to an endotoxin. We also demonstrated that this excitability plasticity is distinctly mediated by translation-independent release of TNF-α via TLR4, and we showed the downstream molecules as MyD88 and TRIF in MG (**Fig. 2**, **Fig. 4** and **Supplementary Fig. 4**). Moreover, the direct injection of bacterial endotoxin into cerebella diminished several types of behaviors of living animals due to functional overconnectivity, which was recovered by both TNF-α inhibition and MG depletion. This was confirmed by FLAIR MR images of inflammation in the cerebellum (**Fig. 5**).

The excitability plasticity of the cerebellar Purkinje neurons is considered to be mediated by M1 MG. Macrophages secrete TNF-α rapidly upon their activation, through non-constitutive pathway^28^. Thus, the mediator release in relatively rapid time scale from activated MG may explain the time course of increase in firing against the exposure to LPS or HKGn (**Fig. 1d,g**) and the translation-independency of the plasticity induction (**Supplementary Fig. 4**). In fact, our results of the increase in TNF-α release at least within 20 minutes from western blotting support this notion (**Fig. 2f,g**). On the other hand, previous studies have shown that glial TNF-α signaling induces trafficking of ionotropic receptors at synapses^12,13^. These trafficking mechanisms of ion-channel receptors underlie protein phosphatase activity. Our results of cerebellar microglia-neuron interaction via TNF-α also suggest the involvement of phosphatase activation for SK channel down-regulation following to TNF receptors in Purkinje neurons (**Fig. 2e**). While TNF-α does not solely target the neuronal membrane, secreted TNF-α is also known to modulate presynaptic transmission via TNF receptors on astrocytes^14,15^. Bergmann glia, a type of astrocytes in the cerebellum, may also promote presynaptic release following TNF-α stimulation. Although our finding regarding the increase in sEPSC frequency (**Fig. 3e,f**, TNF-α) is consistent with this scenario, it’s beyond our scope to address it.

ATP is a gliotransmitter released from neocortical and hippocampal astrocytes upon their Ca^2+^ activity^29,30^. And, intriguingly, astrocytic ATP in the medial prefrontal cortex mediates the antidepressant-like effects through P2X receptors^31^. While several types of cells locate in the cerebellum, none of precisely imaged data regarding the ATP source had been presented. Our ATP imaging (**Fig. 2h-k**) clearly showed that the ATP concentration was increased during the exposure to endotoxin in the ML of the cerebellar cortex, but not in the GL, suggesting the distinct source of ATP in the cerebellum. Cell-level images also suggest that the increase in ATP occurs not in interneurons in the ML, Purkinje cell bodies and dendrites, granule cells, interneurons in the GL or bundles in the white matter (**Supplementary Fig. 5a-g**). And, our pharmacological TNF-inhibition substantially prevented such ATP increase, which suggests the TNF-α secretion from MG is a trigger for the ATP synthesis or its amplification in the ML (**Fig. 2j,k**). Together, the major ATP source in response to the immune triggering is suggested to be Bergman glia or other cells in the ML from our imaging with the new ATP-probe.

Previously, in response to LPS administration ATP has been shown to facilitate the presynaptic release through P2Y receptors in the hippocampal slices^10^ without TNF-α involvement, which is in agreement with our result of the increase in sEPSC frequency by sole ATP administration (**Fig. 3e,f**, ATP). While, in the cerebellum, ATP acts as a diffusible trigger of Ca^2+^ waves through P2Rs in Bergman glial processes^32^, it was shown that such Bergmann glial Ca^2+^ activity itself does not modulate presynaptic release from the glutamatergic excitatory terminals at least on the short time scale^33^. Therefore, ATP-induced Ca^2+^ activity in Bergmann glia and following astrocytic Ca^2+^ dependent messengers may not relate to the increase in sEPSC. Instead, from our results, immune triggered ATP release is suggested to increase the sEPSC frequency of Purkinje cells via P2Rs sustainably (**Fig. 3f**, PPADS+LPS).

Activated MG release not only TNF-α and purines (ATP and GTP), but also other inflammatory cytokines (IL-1β and IL-6), trophic factors (brain-derived neurotrophic factor), and ROS including superoxide and nitric oxide^1–5,8–11,23^. The results of our electrophysiological experiments suggest that ATP and ROS are not involved in the induction of intrinsic excitability increase (**Fig. 2i** and **Fig. 4a**). Rather, considering the reduction of firing frequency of at high ATP concentrations, cerebellar MG could regulate neuronal excitability in bimodal direction through TNF-α and ATP^3^ in a local region. It is also possible that other inflammatory cytokines are involved in this process. Although IL-1β has been shown to prevent the long-term potentiation (LTP) of population spikes in the hippocampal CA1 region^34^, inhibition of IL-1β receptors prevents the maintenance of LTP of population spikes^35^. Interestingly, microglia-specific transcriptomic data suggest changes in gene profile after lesion^6^, and thus, the modulation of neuronal activity would be up to the history of tissue MG. While the relevance of this phenomenon to the late phase of intrinsic plasticity should be examined, our results of western blotting suggest that the protein level of both IL-1β and IL-6 was scarce to detect in response to the acute exposure to LPS in the cerebellum (**Fig. 2f**).

The cerebellum is the principal regulator of motor coordination, timing, and adaptation^36–39^. The vestibular cerebellum for eye movements and reflections, whereas the anterior vermis of the cerebellum has been thought to relate to autonomic nervous system^40,41^. Recently, researchers have started targeting its cognitive functions^26,36,42–44^, and clinical studies have been suggesting involvement of the cerebellum in psychiatric disorders, potentially via dysmetria of thought, manifested by autism spectrum disorders, dyslexia, and schizophrenia^27,44,45^. Akinetic mutism or abulia are also frequently observed after surgical operation of cerebellar astrocytoma or medulloblastoma^46^, suggesting the disruption of connections in regions responsible to speech and motivation through the cerebellar vermis during postoperative inflammation. Histological studies have proved anatomical connection between the cerebellum and neocortex^47,48^. Through evolution, both regions experienced volume expansion correlatively^49^, and thus, such connections may contribute to a wide range of cognitive functions. Here, our present results clarify not only the mechanism of MG-induced hyperexcitability in the cerebellar circuit, but also the psychomotor depressiveness resulting from functional overconnectivity in cerebello-frontal projections (**Fig. 5i**). TNF-α secretion from activated MG appears to be a potential target for suppressing symptoms associated with high cytokine conditions in the inflamed cerebellum that can cause dysfunctional communication (**Supplementary Fig. 14**).

## Acknowledgments

We thank C. Hansel, M. Mitsuyama, P. Hemant, K. Fujita and T. Hirano for invaluable comments on the manuscript and discussions, and the Hakubi-center members for helpful discussions, and H. Tanaka for laboratory support. We thank T. Matsui and T. Matsuda for comments and helpful suggestions regarding rs-*f*MRI analyses and experiments. We thank Y. Itakura and M. Taguchi for assistance with the histology and animal behavior experiments and analyses. We thank Biorbyt Ltd. for offering the CSF1R inhibitor for research purposes. *f*MRI was performed at the Medical Research Support Center, Graduate School of Medicine, Kyoto University, which was supported by Platform for Drug Discovery, Informatics, and Structural Life Science from the Ministry of Education, Culture, Sports, Science and Technology, Japan.

## Author contributions

The funders had no role in study design, data collection and analysis, decision to publish, or preparation of the manuscript; G.O. designed all of the experiments. G.O. (electrophysiology, immunostaining, western blotting, and animal behavior), M.Y. (ATP imaging and transgenic mice generation), M.K. (western blotting and immunostaining), and H.I. (*f*MRI) performed the experiments. G.O. (electrophysiology, ATP imaging, immunostaining, animal behavior, and *f*MRI), M.Y. (ATP imaging), and H.I. (*f*MRI) analysed the data. G.O. and M.K. wrote the manuscript; the authors declare no competing interests.

This work was supported by grants from the Kowa Life Science Foundation, the Japanese Society for Promotion of Science (KAKENHI, Grant-in-Aid for Young Scientists (A) 26710002), the Brain Science Foundation, the Tokyo Biochemical Research Foundation, the Naito Foundation, and the Hakubi-project grant (Kyoto University) (all to G.O.). The funders had no role in study design, data collection and analysis, decision to publish, or preparation of the manuscript. All data is available in the main text or the supplementary materials.

## Additional Information

Reprints and permissions information is available at www.nature.com/reprints. Correspondence and requests for materials should be addressed to G.O.

## Methods

### Animals

Male Sprague-Dawley rats and male GO-ATeam2 (*Related study is in preparation for submission*), TLR2^−/−^, TLR4^−/−^, MyD88^−/−50^, and TRIF^−/−51^ mice (Oriental Bioservice, Inc., Japan) were used for the experiments. Animals were housed (5 animals at maximum in each cage) and maintained under a 12-h light: 12-h dark cycle, at a constant temperature and humidity (20-24°C, 35-55%), with food and water available *ad libitum*. All procedures were performed in accordance with the guidelines of the Animal Care and Use Committees of the local institution and were approved by the Ethical Committee of the local institution. All animal handling and reporting comply with ARRIVE guidelines.

### Patch-clamp recordings

*In vitro* patch-clamp recordings were obtained as described previously^16–19^ Sagittal slices of the cerebellar vermis (250 μm) were prepared from Sprague-Dawley rats (postnatal (P)22-28 days old) after isoflurane anesthesia and decapitation. In some experiments, TLR2^−/−^, TLR4^−/−^, MyD88^−/−^, and TRIF^−/−^ mice (P2-month-old) were used. Slices were cut on a vibratome (Dosaka EM, Japan) using ceramic blades. Subsequently, slices were kept in artificial cerebrospinal fluid (ACSF) containing the following (in mM): 124 NaCl, 5 KCl, 1.25 Na_2_HPO_4_, 2 MgSO_4_, 2 CaCl_2_, 26 NaHCO_3_, and 10 _D_-glucose, bubbled with 95% O_2_ and 5% CO_2_. During cutting, supplemental ingredients (5 mM Na-ascorbate, 2 mM thiourea, and 3 mM Na-pyruvate) were added to the ACSF. After at least 1-h, slices were transferred to a recording chamber superfused with ACSF at near-physiological temperature (32-34°C). The ACSF was supplemented with 100 μM picrotoxin to block GABA_A_ receptors. Patch-clamp recordings were performed under a ×40 water immersion objective lens equipped with a DIC system (DS-Qi2; Nikon) mounted on a microscope (ECLIPSE FN1, Nikon). Recordings were performed in voltage-clamp or current-clamp mode using an EPC-10 amplifier (HEKA Elektronik, Germany). Membrane voltage and current were filtered at 2.9 kHz, digitized at 10 kHz, and acquired using Patchmaster software (HEKA Elektronik). Patch pipettes (borosilicate glass) were filled with a solution containing (in mM): 9 KCl, 10 KOH, 120 K-gluconate, 3.48 MgCl_2_, 10 HEPES, 4 NaCl, 4 Na_2_ATP, 0.4 Na_3_GTP, and 17.5 sucrose (pH 7.25 titrated with 1 M KOH). Membrane voltage was offset for liquid junction potentials (11.7 mV). Somatic patch electrodes had electrode resistances of 2-4 MΩ, while dendritic patch electrodes had electrode resistances of 7-8 MΩ^17^. Hyperpolarizing bias currents (100-400 pA) were injected to stabilize the somatic membrane potential at approximately −75 to −80 mV and to prevent spontaneous spike activity. To obtain the firing frequency in response to different levels of depolarization, we applied 500-ms pulses ranging from 0 to 550 pA every 2 or 3 s, which were increased by 50 pA per step, and counted the number of simple spike-shaped action potentials. Action potential on dendrites were recorded by simultaneous patch-clamping from soma and dendrite. Data were collected at distances of 80-120 μm apart from the soma at secondary and tertiary branches. Due to the lack of voltage-sensitive Na^+^ channels, the amplitude of bAPs attenuates in a distance-dependent manner. Approximate curves obtained from entire recordings including somatic APs are superimposed to the plotting of control (blue line) and apamin (red line) experiments in **Fig. 1i**. To examine the excitability of Purkinje cells from drug-injected cerebella, slices were prepared within 1 h following injection, and recordings were obtained for 1-4 h. For long-term recording, depolarizing current steps (100-400 pA/500 ms) were applied every 20 s to the soma to evoke action potentials. In some experiments, for the conditioning of intrinsic plasticity induction, depolarizing pulses (300-550 pA/100 ms) were applied at 5 Hz for 4 s. We compared firing frequency normalized by 5-min average before 0 min (*i.e.*, −5 - −1 min) to that of 25 to 30 min later. Input resistance was monitored by administering 50-pA hyperpolarizing pulses (50-ms duration) following the depolarization. Data were discarded when the input resistance or the membrane potential had changed more than 20%. All drugs were applied to the bath chamber via the circulation system.

### Electrophysiological data analysis

Data were analyzed using a custom program written in MATLAB (Mathworks). For the analysis of action-potential waveforms, we measured the first action potential evoked by administration of a 200-400-pA depolarizing pulses. Action potential analysis was performed as described previously^16^ (**Supplementary Tables 1** and 2). For the analysis of spontaneous EPSC (sEPSC) events, membrane current was held at −71.7 mV or at −81.7 mV only if the membrane current was jittered, and current was recorded for 1.5 s trials, for at least 180 s in total. Periods of fluctuation were omitted and supplemented by other trials. Then, a Savitzky-Golay filter (*sgolayfilt*) was applied to the recorded currents. The event detection threshold for sEPSCs was set at 4.5 pA. Events were defined as those exceeding four standard deviations during the 10 ms pre-period. We then applied the *fminsearch* function to obtain decay time by a single exponential. If the current trace at the decay period (limited to 19 ms) was poorly fit, the data were excluded. Rise time was regarded as the period spanning 10-90% of the change from peak to basement values. Rise time, half-width, and decay time of sEPSCs were not significantly different against control except for ATP (*data not shown*). Representative sEPSC traces in **Fig. 3b** are the average from 1571, 946, 1264, 1330, 1132, 1079, 663, 276, 1634, and 619 events of control, LPS, PGN, TNF-α, HKEB, HKPA, HKSP, minocycline+LPS, ATP, and PPADS+LPS conditions, respectively. For the cumulative probability in **Fig. 3d** and **3f**, a maximum of 400 sEPSC amplitude and frequency events were collected from each cell for each experiment.

### CSF1R inhibitor treatment

For the sake of pharmacological MG-depletion, the CSF1R inhibitor Ki20227^21,22^ (Biorbyt Ltd., UK) was given to P5-week C57BL/6 mice via the drinking water (100 or 250 mg/L, including 2.5% sucrose) or by oral administration 0.2 mL/day (20 mg/mL dissolved to 10% DMSO and 90% corn oil) for 6-7 days (**Fig. 1j,k** and **Supplementary Fig. 2**). A few mice were given the inhibitor for 11-12 days for electrophysiological experiments. In some experiments, we gave Ki20227 0.2 mL/day (oral administration of 20 mg/mL with 10% DMSO and 90% corn oil) to male juvenile Sprague-Dawley rats (38-62 g body weight) from P19-20 for 5 days. Following reagent administration under specific pathogen free (SPF)-environment, we sacrificed mice for electrophysiological and immunohistochemical experiments, and we conducted behavior tests with rats. No obvious behavioral or health problems were observed during the Ki20227 treatment, except for a reduction of weight in two rats with vulnerability whose data were excluded.

### Immunohistochemistry

Immunostaining was performed as described^16^, with some modifications. After perfusion fixation of the control and Ki20227-treated mice with 4% paraformaldehyde (PFA), brains were kept in PFA for 2 days at 4°C. After submersion in phosphate buffered saline (PBS) containing 30% sucrose for 2-4 days, 50-μm cryosections were collected in water. Sections were heated in HistoVT (Nacalai Tesque, Japan) to 80°C for 30 min, rinsed in TBS, and incubated in blocking solution (TBS containing 10% normal goat serum (NGS) and 0.5% Triton) for 1 h at 20-24°C. Sections were then incubated with fluorescently conjugated-primary antibodies against Iba1 (rabbit anti-Iba1:red fluorescent probe 635, 1:200 [2.5 μg/mL]; Wako) and Calbindin (rabbit anti Calbindin:Alexa Fluor^®^ 488, 1:100 [5 μg/mL]; Abcam) at 4°C for 48 h. Subsequently, sections were rinsed three times in PBS for 5 min each and were mounted on glass slides and cover-slipped. Fluorescence images were obtained using a Zeiss LSM 780, Olympus FV1000 or Olympus FV 3000 confocal laser-scanning microscope equipped with Plan-Apochromat 20×/0.8 and 10×/0.4 lenses. Emission wave length for imaging was 488 and 639 nm, and the fluorescence was filtered using 640nm low-pass and 490-555-nm band-pass filters. Individual images were taken under a fluorescence microscope (FV 3000), and the merged-images are shown as whole cerebellar images in **Supplementary Fig. 2**. For counting the number of MG in **Fig. 1k**, arbitral parts of the cerebellum were imaged (489.5 ×489.5 μm) with z-stacks of 24-30 images at every 1 μm, and the density of MG were calculated from 43-54 regions of interest (ROIs) (from 2 mice per group), excluding white matter.

### Western blotting

Cerebellar slices from Sprague-Dawley rats were prepared as described above. After recovery, brain slices were incubated in normal ACSF or LPS-containing ACSF for 0, 20, or 60 min at near-physiological temperature. Supernatants were concentrated with Amicon Ultra-15 Centrifugal Filter Units (EMD Millipore) and subjected to immunoblotting analysis using anti-rat TNF-α (BMS175; eBioscience), anti-rat IL-1β (AF506; R&D) and anti-rat IL-1β (AF-501-NA; R&D). Intensity was quantified using Multi Gauge ver.3.2 (Fujifilm). For a control experiment, we used rat macrophage culture (NR8383 [AgC11x3A, NR8383.1], ATCC) (*data not shown*).

### Reagents

Apamin, apocynin, cyclosporin A, C34^52^, C87^53^, minocycline hydroxide^54^, PPADS tetrasodium salt, picrotoxin, NBQX disodium salt, anisomycin, and cycloheximide (Tocris); LPS from *E coli O26* or *O111* (Wako); Ki20227^22^ (IC_50_= 2 nM to M-CSF receptor; 451 nM to c-Kit), which is a more potent and specific inhibitor to CSF-1R than PLX3397^55^ (IC_50_ = 20 nM to M-CSF receptor; 10 nM to c-Kit); okadaic acid (AdipoGen); ATP (Adenosine 5’-triphosphate disodium salt), BAPTA (Sigma-Aldrich); carrier-free recombinant rat TNF-α (R&D Systems); heat-killed bacteria (HKEB, *E.coli 0111:B4;* HKPA, *P. aeruginosa;* HKSP, *S. pneumoniae*), and peptidoglycan (PGN, peptidoglycan from *E. coli 0111:B4*) were purchased from InvivoGen, and 10^10^ freeze-dried cells were diluted to 10^7^ or 10^9^ cells/mL in sterile, endotoxin-free water.

### ATP imaging

An ATP probe (GO-ATeam2) was developed for use in conjunction with green and orange fluorescent proteins as a fluorescence/Förster resonance energy transfer (FRET) pair^56^. We generated GO-ATeam2 transgenic mice to monitor the ATP concentration in living animals. Briefly, we employed a knock-in strategy targeting the Rosa26 locus and the CAG promoter to regulate transcription. We used the GeneArt Seamless Recombination System (Thermo Fisher Scientific) to create GO-ATeam2 knock-in mice. The targeting vector was induced into G4 ES cells with electroporation. The constructs harbored by the ES clones underwent homologous recombination, which was confirmed by Southern blot analysis using appropriate probes (provided by K. Hoshino and T. Kaisho, Osaka University), PCR, and qPCR. Male chimeras derived from each ES cell line were bred with C57BL/6J females, yielding heterozygous F1 offspring (C57BL/6J × 129 background). P2-3 weeks GO-ATeam2 mice were decapitated after inhalation of 2% isoflurane, following which whole brains were isolated. The cerebellum was placed in cooled ACSF solution as described for the electrophysiological experiments. Air stones (#180; Ibuki) were used for aeration (95% O_2_ and 5% CO_2_). Sagittal slices were cut to a thickness of 300 μm using a vibratome (VT 1000S; Leica), following which they were maintained for at least 30 min at room temperature in ACSF solution. FRET imaging was performed on a two-photon microscope (TCS SP8; Leica). The imaging chamber was set on the stage of the microscope with flowing ACSF solution bubbled with 95% O_2_ and 5% CO_2_. LPS (final concentration 12 μg/mL) or a mixture of HKEB and HKPA (final 10^7^ cells/mL, for each) were added to the ACSF. Exciting light (920 nm, 25 W under objective lens) was applied, and the molecular layer (ML) and granule cell layer (GL) of the cerebellum were scanned every 2 min. Images were obtained from individual locations (scanned area size, 550 × 550 μm). We used BP525/50 filters for emission, and DM560 and D585/40 filters for excitation fluorescence separation. IMD images and quantifying images were developed from the fluorescence images using MetaMorph software (Roper Scientific, Trenton, NJ). FRET signals at the chosen ROI (whose shape depended on the target area, **Supplementary Fig. 5h**) in the ML and GL were averaged, and the obtained ratio was applied to the following equation: FRET ratio = 1.52 × [ATP]^1.7 / ([ATP]^1.7 + 2.22) + 0.44, and then, ATP concentration was calculated. The coefficients were carefully determined based on two methods. First, mouse embryonic fibroblasts were obtained from GO-ATeam2 mice. After piercing the fibroblast membrane, we applied different concentrations of ATP, monitored the FRET ratio, and determined coefficients based on the function for fitting ATP concentration to the FRET ratio. Second, we performed a luciferase assay of ATP concentration (Tissue ATP assay kit; TOYO B-NET) in fertilized eggs from GO-ATeam2 mice. Mice were injected with ATP synthetase inhibitors (2DG plus antimycin A), and the time course of the FRET ratio was monitored. Coefficients obtained by the two methods were substantially identical, indicating that they were appropriate for use in the present study. Our ATP imaging with transgenic mice expressing ATP probes monitored the ATP concentration in the cytosol, insomuch as no specific promoters were tied to the transgene construct. Control fluorescence images were obtained prior to drug exposure (n = 9 in ML, and n = 10 in GL) and used for subtraction of the background signal. Following endotoxin exposure, remnants on the tubing and chamber were cleaned with 80% ethanol for at least 5 min. To visualize the time courses of ΔATP, baseline ATP was set to zero (−6 to −2 min).

### Drug injection

Rats received an injection of 0.20-0.30 μL of PBS, LPS (1 mg/mL), a mixture of heat-killed Gram-negative bacteria (HKEB and HKPA, 10^9^ cells/mL for each), a TNF-α inhibitor (C87, 2-4 mM), TNF-α (20 μg/mL) and ATP (20 mM) into the vermis or right hemisphere of the cerebellum. Drugs were injected under anesthesia with 0.9% ketamine (Daiichi Sankyo Co., Ltd.) and 0.2% xylazine (Bayer AG) (*i.p.*, 5.3 ± 0.5 μL/g of body weight, as mean ± std.). We started surgery after animals’ breath and pulse were stabilized and the extent of anesthesia was enough, without corneal reflection, touch, and pinch responses. For vermal injection into cerebella, rats were fixed to the stereotaxic apparatus and a small hole (300 μm radius) was drilled in the skull (2.5 mm posterior to lambda), following which a microsyringe was inserted forward to the anterior lobule (3.0-3.5 mm depth at 86°, lobule II-IV)^57^. For injection into the hemispheres, a hole was made at 3.0-3.5 mm posterior to the lambda and around 1.5 mm lateral, putting forward to 4.0 mm depth with an angle of 60-65°. The wound was sealed with Spongel (Astellas Pharma Inc.), after which the animal was allowed to rest in a clean cage. A heat-plate was used to maintain body temperature, if necessary. Rats woke up approximately 40 min after injection of the ketamine/xylazine. We handled animals very carefully and treated them with as little discomfort as possible. After at least 2 h of recovery, we confirmed that the effects of the anesthesia had dissipated and that no paralysis or seizures had occurred, following which the behavior test battery (open field test, social interaction test, forced swim test, marble burying test, retention test on balance beam, and gait analysis) was initiated. Forced swim test was conducted last in the battery, to avoid the effect of fear conditioning itself. Gait analysis and retention test on balance bar after recovery revealed no obvious ataxia and motor discoordination (**Supplementary Fig. 11a,b** and **Supplementary Fig. 13f**).

### Open field test

We monitored the spatial exploratory behavior of Sprague-Dawley rats (P22-26 days) weighing 52-86 g and microglia-depleted rats (P24-25 days). After habituation in the experimental room (1 h), rats were individually placed on the center of the Plexiglas open field arena (72 × 72 cm^2^, 30 cm high white walls with black floor), following the operation. We monitored the behavior of freely moving rats for 30 min using a video camera. The distance travelled, resting period, moving period, and mean speed were compared among the groups. The arena and surrounding walls were cleaned and deodorized with H_2_O and 70% EtOH before each session. Exploration behavior was quantified using Smart 3.0 software (Panlab Harvard Apparatus). The resting state was defined at that during which moving speed fell below the threshold of 2.5 cm/s. A total of 13, 11, 11, 11, 12, 15, 12, 16, and 11 animals were included in the non-conditioned (NC), PBS, LPS, heat-killed Gram-negative bacteria mixture (HKGn; HKEB+HKPA), HKGn+C87, TNF-α, ATP, MG-depleted (dMG)+LPS, and LPS to hemispheres conditions, respectively.

### Social interaction

The sociability of rats was tested in a Plexiglas open-field arena (72 × 72 cm^2^, 30 cm high) with small circular wire cages (15 cm in diameter, 20 cm high) in two corners. Animals were allowed to explore the arena for 3 min to determine the baseline of exploratory behavior against the novel subjects without social targets. All movements were recorded with video-tracking. We defined a preferred corner area as one area (30 × 36 cm^2^ square around a wire cage) where the animal spent more time during the baseline period. Next, a sibling rat was placed into one cage that was less preferred during the baseline period, and movements were monitored for another three minutes. In this social approach model, time spent in the interaction and overall locomotion were compared by an examiner in a blinded experimental condition. Sociability index was calculated by dividing the time difference between time spent in the interaction zone and in other areas with and without a sibling by total time (**Fig. 5c** and **Supplementary Fig. 13c**), as following; Sociability index = ((Ta^test^ − Tā^test^) − (Ta^baseline^ − Tā^baseline^)) / T^total^, where T as the resident time (sec), Ta^test^ as time spent in the area where the sibling caged in the test period, Tā^test^ as time spent out of the area where the sibling is caged, and T^total^ as 180 seconds. Ta^baseline^ and Tā^baselme^ are those in the baseline period. In Fig. 5c, abbreviations are given in the equation of sociability index. A total of 15, 15, 12, 13, 16, 13, and 16 animals were included in the PBS, LPS, HKGn, HKGn+C87, TNF-α, ATP, and dMG+LPS conditions, respectively.

### Forced swim

FS test were conducted in a 5 L plastic beaker filled with 4.3 L of water (16.5 cm in diameter, 20-cm-depth, 24.1 ± 0.2°C). Rats were tested swimming for 8 min and video-recorded. Total duration of immobility, each of which is more than 2 s, in the last 4 min was measured in a blinded condition. Total of 15, 15, 16, 14, 16, 13, and 16 animals were included in the PBS, LPS, HKGn, HKGn+C87, TNF-α, ATP, and dMG+LPS conditions, respectively.

### Marble burying

Marble burying (MB) is a test for stereotyped repetitive behaviors in rodents analogous to those observed in autistic phenotypes^58^. MB tests were conducted in a testing cage (26 × 18.2 cm^2^, 13-cm-high). Bedding tips as wood shavings were covered to a depth of 3 cm. 20 glass marbles (17 mm in diameter) were aligned equidistantly in four rows of five marbles each. Spaces (4-5 cm width) were made for placing animals. At the end of the 20-min test period, rats were carefully removed from the cages. The marble burying score was defined as the following: 1 for marbles covered >50% with bedding, 0.5 for marbles covered ~50% with bedding, and 0 for marbles less covered. A total of 24, 16, 15, 16, 13, 15, 13, and 14 animals were included in the NC, PBS, LPS, HKGn, HKGn+C87, TNF-α, ATP, and dMG+LPS conditions, respectively.

### Retention test on balance beam

Time in seconds was measured while rats remained on a wooden balance beam with 25 mm diameter placed at a height of ~11 cm. Measurement time is 5 minutes after animals became calm on the beam. Trials in which animals escaped were not analyzed. A total of 12, 10, 12, 10, 16, 11, 13, and 12 animals were included in the PBS, LPS, HKGn, HKGn+C87, TNF-α, ATP, dMG+LPS, and LPS to hemispheres conditions, respectively.

### Gait analysis

Animals (P23-26) were placed at the end of one directional passageway (8.5 cm in width and 30 cm in length) with a transparent floor at a height of 13 cm, and they were allowed to walk straight forward while being recorded with a micro video camera from below. Each animal was tested three times for walking. Centers of paw positions (forepaws and hind paws) were measured, and three or four strides lengths for each trial were collected from 6, 6, 6, 7, 6, and 7 rats of PBS, LPS, HKGn, HKGn+C87, dMG+LPS, and LPS to hemispheres conditions.

### MR imaging and data analyses

We used anaesthetized Sprague-Dawley rats as described in the open field test, with three experimental groups of non-conditioned (NC, n = 13), HKGn-injected (HKGn, n = 12), and HKGn+C87-injected (HKGn+C87, n = 14) rats. After a 2-h recovery period from the ketamine and xylazine anesthesia as described above, rats inhaled isoflurane (2% for induction: 1% for MR imaging (MRI)) in a mixture of 66% air and 34% oxygen at 1.5 L/min with ventilation, were stabilized by head-holding in a plastic tube, and were monitored for respiratory rate (52-109 breaths/min) and body temperature (30-34°C). Rats were scanned with a 7.0 T MRI scanner (Bruker BioSpin) with a quadrature transmit-receive volume coil (35 mm inner diameter). Shimming was performed in a 20 × 15 × 10 mm^3^ region by mean of a local MapShim protocol using a previously acquired field map. Blood-oxygen-level dependent (BOLD) resting state- (rs-)*f*MRI time series were obtained with a single-shot gradient-echo planar imaging (EPI) sequence (repetition time [TR]/echo time [TE] = 1.0 s/9 ms; flip angle, 60°; matrix size, 80 × 64; field of view [FOV], 2.5 × 2.0 cm^2^; 12 coronal slices from top to bottom; slice thickness, 1 mm; slice gap 0 mm) for 6-8 min with a total 360-480 volumes. Following the EPI sequence twice, high-resolution anatomical images for each experimental animal were obtained using a 2D multi-slice T_2_-weighted (T2W) fast-spin echo sequence (RARE) (TR/TE = 3.0 s/36 ms; matrix size, 240 × 192; FOV, 2.5 × 2.0 cm^2^; 24 coronal slices; slice thickness, 0.50 mm; slice gap 0 mm; with fat suppression by frequency selective pre-saturation) under 2.0% isoflurane inhalation. To image the region of inflammation, we used a fluid attenuation inversion recovery (FLAIR) sequence, which suppresses cerebrospinal fluid effects on the image (inversion time [TI] = 2.5 s; TR/TE = 10 s/36 ms; matrix size, 240 × 192; FOV, 2.5 × 2.0 cm^2^; 24 coronal slices; slice thickness, 0.50 mm; slice gap 0 mm; with fat suppression by frequency selective pre-saturation). Image data from three experiments of NC, HKGn-injected and HKGn+C87-injected were analyzed with SPM12 (http://www.fil.ion.ucl.ac.uk/spm), FSL (https://fsl.fmrib.ox.ac.uk/fsl/fslwiki/FSL), and in-house software written with MATLAB (MathWorks). We pre-processed imaging data as described previously^59,60^. First, the EPIs were realigned and co-registered to a template brain using anatomical images. Owing to the lack in open source anatomical brain images in young adult rats, we used a representative T2W anatomical image (P22) as a template brain. Co-registered functional EPIs were normalized to the template and transformed to a 151 × 91 × 81 matrix (with spatial resolution of 0.20 × 0.20 × 0.20 mm^3^) and smoothed with a Gaussian kernel (full width at half maximum [FWHM], 0.7 mm). We manually omitted data with motion artefacts (26 scans in total). Imaging data were temporally zero-phase band-pass filtered to retain low-frequency components (0.01-0.10 Hz) by using the *filtfilt* Matlab function. For a given time series, seed ROIs (0.8 × 0.8 × 0.6 mm^3^) in the anterior lobe of the cerebellar vermis (CblVm), cerebellar hemispheres (CblHs), cerebellar dentate nuclei (CblNc), dorsal hippocampi (Hpc), primary visual cortices (V1), sensory cortices (S1), motor cortices (M1), cingulate cortices (Cg), medial prefrontal cortices (mPf), centro-medial thalamus (cmThl), and posterior thalami (pThl) were selected, and the signals in the seed were averaged. For CblHs, CblNc, Hpc, V1, S1, M1, Cg, mPf, and pThl, seeds were applied in both right and left hemispheres. Individual correlation maps (r map at the zeroth lag) were computed by cross-correlation against the mean seed-signal to signals of all the other voxels. Then, correlation maps were transformed to normally distributed z scores by Fisher’s r-to-z transformation. Z-transformation was used to reflect the strength of spontaneous correlations more linearly at high r values. The resulting group maps were thresholded at |r| > 0.1, followed by a cluster-level multiple comparison correction at a significance level of p < 0.001 of one-sample t-test with Kα > 29 voxels (**Fig. 5g**). Seed-seed correlation matrices were calculated from all pairs among brain regions for each subject. The correlation matrix was z-transformed. One-sample *t-test* matrix across experiments were thresholded at p < 0.05 and were corrected by Benjamini-Hochberg (BH) procedure to avoid the incorrect rejection of a true null hypothesis (a type I error) with a false discovery rate at q = 0.05. Mean seed seed correlation z-matrix was filtered by the T-matrix from BH procedure and variance corrected color maps were shown (**Fig. 5h**).

For the group independent component analysis (ICA) (**Supplementary Fig. 15**), we used the MELODIC toolbox in the FSL platform^61,62^. Group ICA was done on the experimental groups (NC, HKGn, and HKGn+C87) to estimate a common set of components for all three cohorts, using multi-session temporal concatenation, and we then extracted 40 components from the pre-processed data described above. The set of spatial maps from the three-cohort-average analysis was used to generate subject-specific versions of the spatial maps, using dual regression. We then generated average spatial maps for each group by one-sample t-test using the randomize function of FSL^63^. The resulting maps were corrected for multiple comparisons using threshold-free cluster enhancement^64^. These representative maps are color-coded as a value of 1-p with a threshold of p < 0.01.

### Statistics

All data are presented as mean ± SEM unless otherwise stated. The summary of statistic is tabulated in **Supplementary Table 3**. Two-sided Mann-Whitney *U*-tests were used to compare data between two independent groups, except for the following: in **Fig. 1k**, **Fig. 5b-e**, **Supplementary Fig. 11** and **Supplementary Fig. 13**, we used the Kruskal-Wallis test with multiple comparison test, with Bonferroni method and Fisher’s least significant difference procedure, respectively. In **Fig. 1n** and **Fig. 4c-g**, we used the Wilcoxon signed-rank test between the mean normalized firing frequency at −1 to −5 min and at +25 to +30 min. In **Fig. 2i** and **2k**, we used the Wilcoxon signed-rank test between the mean ΔATP at −6 to −2 min and at +20 to +28 min. In **Supplementary Fig. 12b**, we used the unpaired Student’s *t*-test (two-tailed, unequal distribution). p < 0.05 was considered statistically significant, unless otherwise stated. All boxplot graphs show interquartile range with centered bars as median. Overlapping red marks on boxes represent mean ± SEM. Other thresholds are provided for each relevant comparison. Outliers in **Fig. 5b** (Total distances (n=3), Mean speed (n=1), and FS immobility (n=1)) were excluded under assumption of normality by Grubb’s test.

## Reporting Summary

Further information on experimental design is available in the Nature Research Reporting Summary linked to this article.

## Data availability statement

Source Data that support the findings of this study are available in the online version of the paper and from the corresponding author upon reasonable request.

## Code availability

Custom Matlab code for analyses are available from the corresponding author upon individual request.

---

**Supplementary Table 1.**
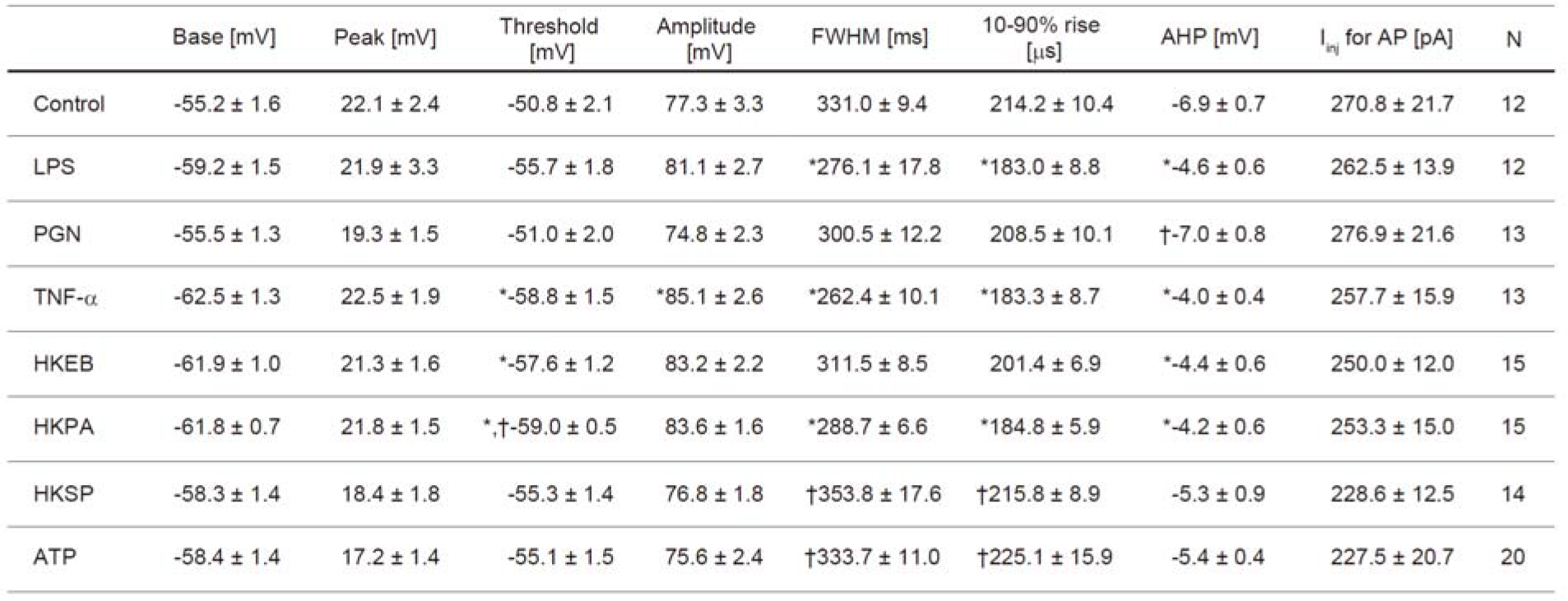
Action potential waveform parameters of rat Purkinje neurons in response to immune-related reagents. Recordings were obtained under current-clamp. Basement voltage, action potential (AP) peak, AP threshold, AP amplitude to the peak, AP width as full width at half maximum (FWHM), 10-90% rise time, after hyperpolarization (AHP), and injected current are shown with the corresponding number of cells. APs were generated by a depolarization current administered at the soma. Data are shown as mean ± SEM. p-values were obtained using two-sided Mann-Whitney *U*-test between control or LPS and other factors. Significant differences are marked with * and ^†^, respectively. ATP, adenosine triphosphate; HKEB, heat-killed *Escherichia coli;* HKPA, heat-killed *Pseudomonas aeruginosa;* HKSP, heat-killed *Streptococcus pneumoniae;* LPS, lipopolysaccharide; PGN, peptidoglycan; TNF-α, tumor necrosis factor alpha.

**Supplementary Table 2.** Action potential properties of microglia-depleted and immune-related molecule-deficient mouse Purkinje neurons. Recordings were obtained under current-clamp. Basement voltage, action potential (AP) peak, AP threshold, AP amplitude to the peak, AP width as full width at half maximum (FWHM), 10-90% rise time, after hyperpolarization (AHP), and injected current are shown with the corresponding number of cells. APs were generated by a depolarization current administered at the soma. Data are shown as mean ± SEM. p-values were obtained using two-sided Mann-Whitney *U*-test between control or LPS and other factors. Significant differences compared among each group are marked with *. MG depleted, microglia-depleted (dMG); HKGn, heat-killed Gram-negative bacteria mixture; LPS, lipopolysaccharide; MyD88 KO, Myeloid differentiation primary response gene 88 knockout mice; TLR2/4 KO, Toll-like receptor 2/4 knockout mice; TRIF KO, TIR-domain-containing adapter inducing interferonβ knockout mice.

| | Base [mV] | Peak [mV] | Threshold [mV] | Amplitude [mV] | FWHM [ms] | 10-90% rise [ms] | AHP [mV] | $I_{inj}$ for AP [pA] | N |
| --- | --- | --- | --- | --- | --- | --- | --- | --- | --- |
| MG-depleted | -57.5 ± 1.9 | 20.5 ± 1.4 | -53.1 ± 2.4 | 78.0 ± 2.5 | 271.3 ± 6.8 | 224.6 ± 9.9 | -7.0 ± 1.1 | 304.2 ± 45.4 | 12 |
| dMG + LPS | -61.0 ± 1.8 | 22.6 ± 1.9 | -57.0 ± 2.1 | 83.6 ± 2.4 | 262.3 ± 5.2 | 195.2 ± 10.2 | -6.9 ± 0.7 | 236.7 ± 27.8 | 15 |
| dMG + HKGn | -57.8 ± 1.2 | 21.3 ± 1.7 | -52.9 ± 1.7 | 79.0 ± 2.6 | 280.2 ± 9.6 | 232.7 ± 17.4 | -7.4 ± 0.7 | 258.8 ± 25.8 | 17 |
| TLR2 KO | -58.2 ± 1.8 | 14.4 ± 2.4 | -53.6 ± 2.5 | 72.6 ± 3.3 | 261.5 ± 15.8 | 238.8 ± 24.1 | -7.4 ± 0.9 | 238.9 ± 43.9 | 9 |
| TLR2 KO + LPS | *-64.3 ± 1.0 | 17.3 ± 2.2 | *-60.7 ± 0.9 | *81.6 ± 2.7 | 277.8 ± 10.3 | 189.5 ± 7.1 | -5.5 ± 0.4 | 191.7 ± 5.6 | 12 |
| TLR4 KO | -55.7 ± 2.1 | 14.1 ± 2.1 | -49.4 ± 2.7 | 69.8 ± 2.4 | 291.5 ± 11.5 | 223.6 ± 20.0 | -7.9 ± 0.9 | 278.6 ± 1.5 | 14 |
| TLR4 KO + LPS | -56.8 ± 1.8 | 13.0 ± 2.0 | -51.7 ± 2.4 | 69.8 ± 3.1 | 289.1 ± 12.2 | 236.8 ± 21.0 | -7.2 ± 0.9 | 250.0 ± 24.4 | 15 |
| TLR4 KO + HKGn | -60.9 ± 1.6 | 8.7 ± 1.7 | -56.9 ± 2.1 | 69.6 ± 2.5 | 316.7 ± 11.0 | 218.1 ± 11.3 | -6.1 ± 0.7 | 220.0 ± 17.5 | 15 |
| MyD88 KO | -59.4 ± 1.7 | 16.9 ± 1.5 | -53.9 ± 2.4 | 76.4 ± 2.4 | 240.7 ± 9.0 | 192.6 ± 11.7 | -7.1 ± 0.5 | 281.8 ± 23.6 | 12 |
| MyD88 KO + LPS | -61.4 ± 2.2 | 17.9 ± 2.8 | -56.0 ± 2.4 | 79.4 ± 4.4 | 254.9 ± 9.0 | 198.3 ± 12.6 | -6.8 ± 0.7 | 277.3 ± 23.7 | 11 |
| TRIF KO | -59.4 ± 1.3 | 21.4 ± 1.7 | -54.5 ± 2.0 | 80.8 ± 1.8 | 265.8 ± 9.2 | 188.0 ± 9.1 | -6.5 ± 0.7 | 253.3 ± 19.2 | 15 |
| TRIF KO + LPS | -59.9 ± 1.4 | 20.3 ± 1.5 | -54.6 ± 1.9 | 80.2 ± 1.9 | 264.8 ± 8.2 | 200.6 ± 9.2 | -6.4 ± 0.8 | 276.5 ± 20.2 | 17 |

**Supplementary Table 3.** Summary of the statistics. The statistical methods, numbers of data, statistic results, and statistic values in each figure are tabulated (1-5).

| Figure number | Statistic method | The number of data | Statistic results | Statistic values |
| --- | --- | --- | --- | --- |
| 1b<br>LPS vs Control | Two-tailed<br>Mann-Whitney<br>*U*-test | Control, n = 14<br>LPS, n = 12<br>{50, 100, ... , 550 pA} | p = {0.1331 0.0500 0.0188<br>0.0152 0.0167 0.0179 0.0179<br>0.0117 0.0059 0.0039 0.0052} | U = {70.0 48.5 38.5<br>36.5 37.0 37.5 37.5<br>34.5 30.0 27.5 25.5} |
| 1d<br>LPS vs Control | Two-tailed<br>Mann-Whitney<br>*U*-test | Control, n = 8<br>LPS, n = 8 | p = 0.00062 | U = 2 |
| 1e<br>HKEB vs Control | Two-tailed<br>Mann-Whitney<br>*U*-test | HKEB, n = 15<br>{50, 100, ... , 550 pA} | p = {0.0453 0.0015 0.0040<br>0.0066 0.0077 0.0144 0.0230<br>0.0323 0.0072 0.0015 0.0008} | U = {62.0 20.0 24.0<br>27.5 28.5 33.5 37.5<br>40.5 28.0 17.0 13.0} |
| 1e<br>HKPA vs Control | Two-tailed<br>Mann-Whitney<br>*U*-test | HKPA, n = 15<br>{50, 100, ... , 550 pA} | p = {0.0048 0.0024 0.0019<br>0.0031 0.0045 0.0055 0.0063<br>0.0059 0.0012 0.0003 0.0004} | U = {41.0 24.0 19.0<br>22.0 24.5 26.0 27.0<br>26.5 15.5 6.5 8.0} |
| 1e<br>HKSP vs Control | Two-tailed<br>Mann-Whitney<br>*U*-test | HKSP, n = 14<br>{50, 100, ... , 550 pA} | p = {NaN 0.1255 0.1171<br>0.0718 0.0687 0.1024 0.3456<br>0.6287 0.4616 0.2691 0.1677} | U = {98.0 67.0 64.0<br>58.5 58.0 62.0 77.0<br>87.0 81.5 73.5 67.5} |
| 1g<br>HKEB vs Control | Two-tailed<br>Mann-Whitney<br>*U*-test | HKEB, n = 6 | p = 0.0080 | U = 4 |
| 1g<br>HKPA vs Control | Two-tailed<br>Mann-Whitney<br>*U*-test | HKPA, n = 6 | p = 0.0127 | U = 5 |
| 1g<br>HKSP vs Control | Two-tailed<br>Mann-Whitney<br>*U*-test | HKSP, n = 7 | p = 0.1419 | U = 12 |
| 1i<br>LPS vs Control | Two-tailed<br>Mann-Whitney<br>*U*-test | Control, n = 28<br>LPS, n = 5 | p = 0.0328 | U = 113 |
| 1i<br>Apamin vs<br>Control | Two-tailed<br>Mann-Whitney<br>*U*-test | Control, n = 28<br>Apamin, n = 9 | p = 0.0014 | U = 217 |
| 1k | Kruskal-Wallis test<br>with Bonferroni<br>method | Control, n = 54<br>100µg/ml, n = 48<br>250µg/ml, n = 44<br>20mg/ml o.a., n = 43 | p = 0, for Kruskal-Wallis test<br>p = 4.7*10<sup>-10</sup>, for Ctrl vs 100µg/ml<br>p = 8.4*10<sup>-11</sup>, for Ctrl vs 250µg/ml<br>p = 6.1*10<sup>-35</sup>, for Ctrl vs 20mg/ml<br>p = 1.0, for 100µg/ml vs 250µg/ml<br>p = 1.2*10<sup>-8</sup>, for 100µg/ml vs 20mg/ml<br>p = 2.4*10<sup>-7</sup>, for 250µg/ml vs 20mg/ml | Kruskal-Wallis statistics<br>= 157.66 |
| 1l<br>dMG vs Control | Two-tailed<br>Mann-Whitney<br>*U*-test | dMG, n = 12<br>{50, 100, ... , 550 pA} | p = {0.1331 0.3962 0.9777<br>0.5881 0.4005 0.5691 0.8368<br>0.8772 0.9590 0.5198 0.2363} | U = {97.0 96.5 83.0<br>73.5 67.5 72.5 79.5<br>80.5 85.5 97.0 87.5} |
| 1l<br>dMG+LPS vs<br>Control | Two-tailed<br>Mann-Whitney<br>*U*-test | dMG+LPS, n = 14<br>{50, 100, ... , 550 pA} | p = {0.0684 0.3133 0.1784<br>0.3375 0.4077 0.5927 0.5595<br>0.4515 0.2853 0.1519 0.0847} | U = {84.0 86.5 77.5<br>85.0 87.5 93.5 92.5<br>89.0 82.5 75.0 69.0} |
| 1l<br>dMG+HKGn vs<br>Control | Two-tailed<br>Mann-Whitney<br>*U*-test | dMG+HKGn, n = 17<br>{50, 100, ... , 550 pA} | p = {NaN 0.5342 0.7944<br>0.7917 0.8260 0.9524 0.8580<br>0.7356 0.3819 0.1470 0.0619} | U = {119.0 105.5 125.5<br>126.0 125.0 121.0 114.0<br>110.0 96.5 82.0 71.5} |
| 1n<br>dMG+5Hz-depolariza-<br>tion | Wilcoxon<br>signed-rank test | dMG+5Hz-depol.,<br>n = 6 | p = 0.0313 | signed-rank test statistics<br>= 0 |
| 1n<br>dMG+LPS | Wilcoxon<br>signed-rank test | dMG+LPS, n = 8 | p = 0.0234, for reduction | signed-rank test statistics<br>= 2 |
| 1n<br>dMG+HKGn | Wilcoxon<br>signed-rank test | dMG+HKGn, n = 6 | p = 0.9375 | signed-rank test statistics<br>= 10 |

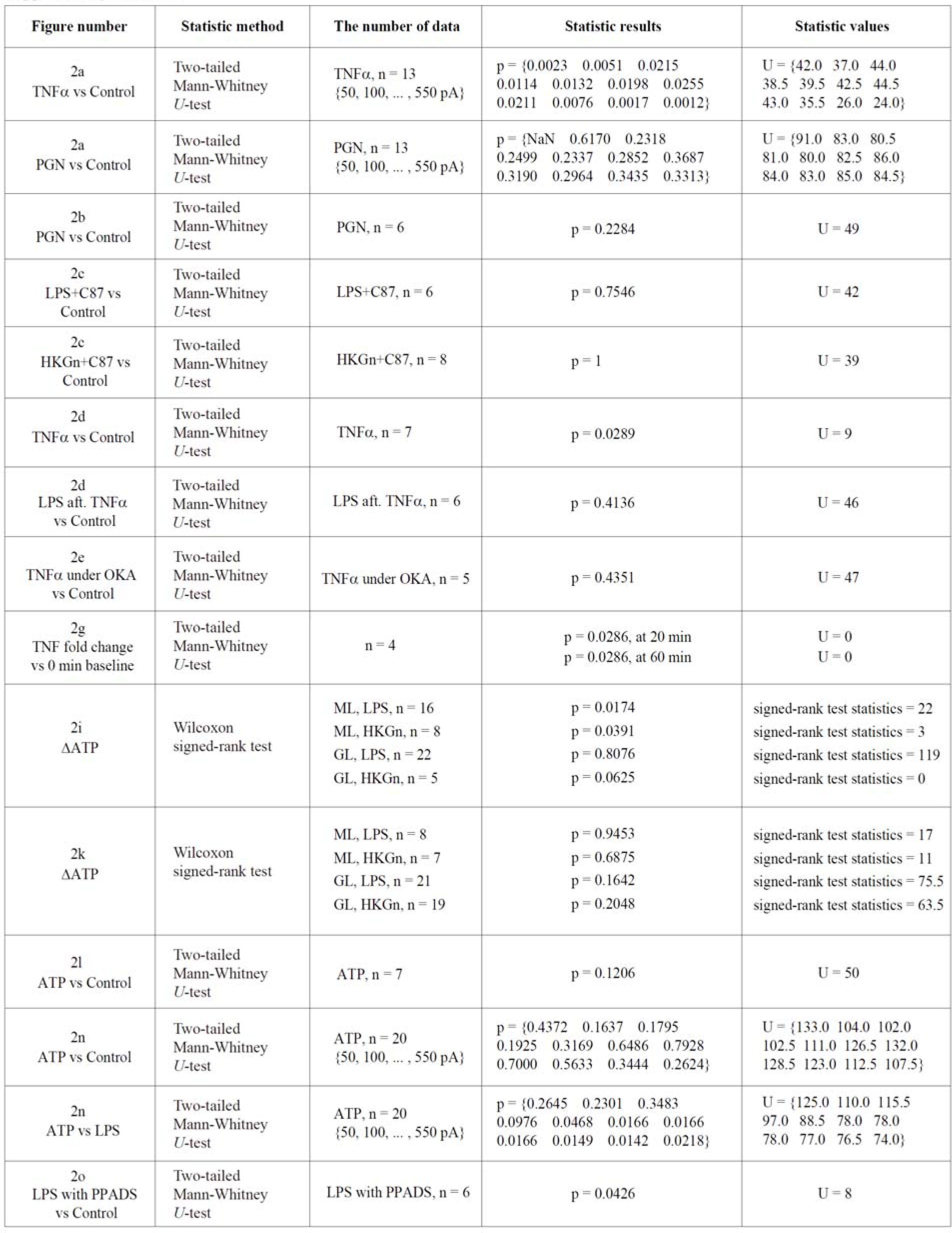

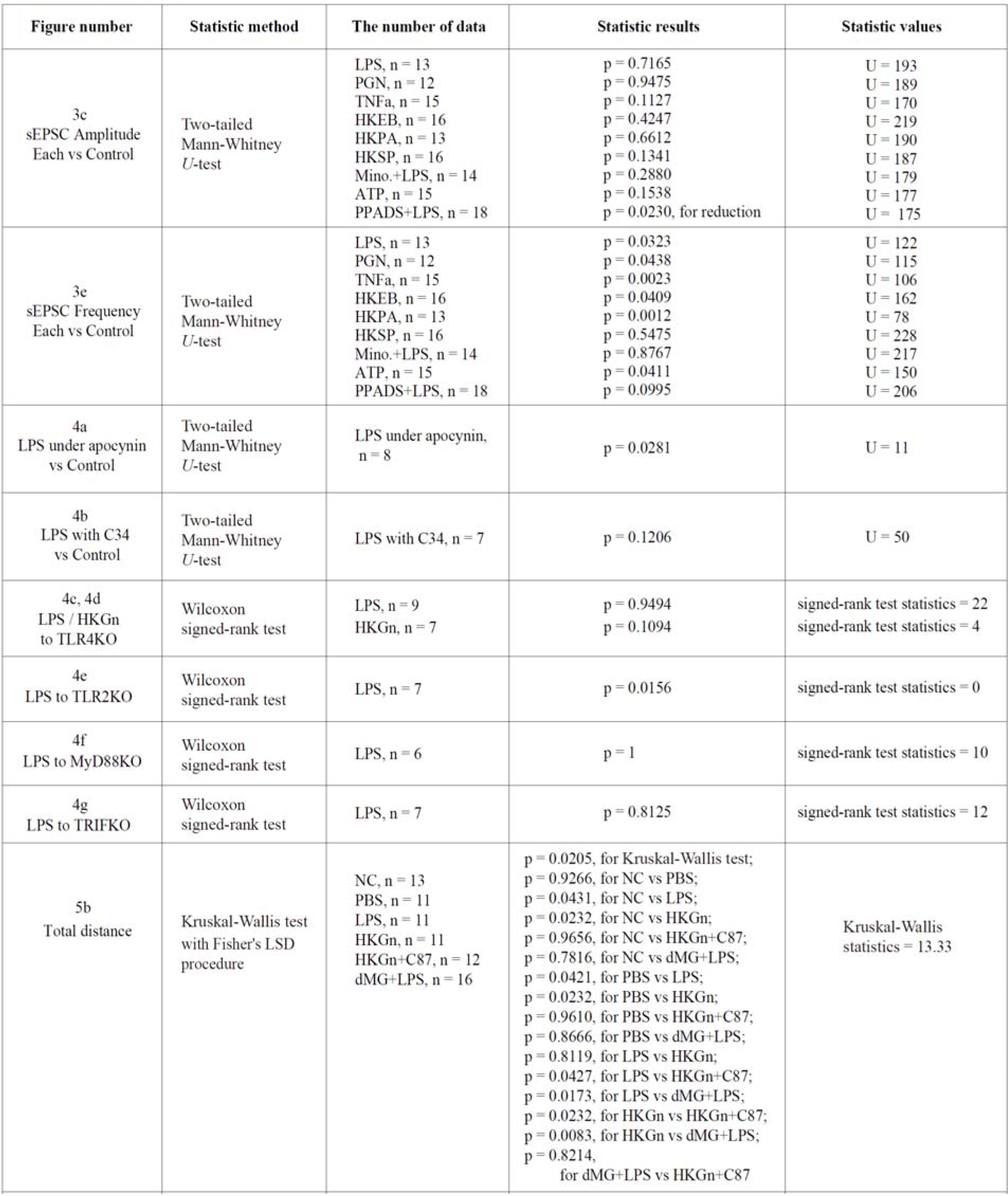

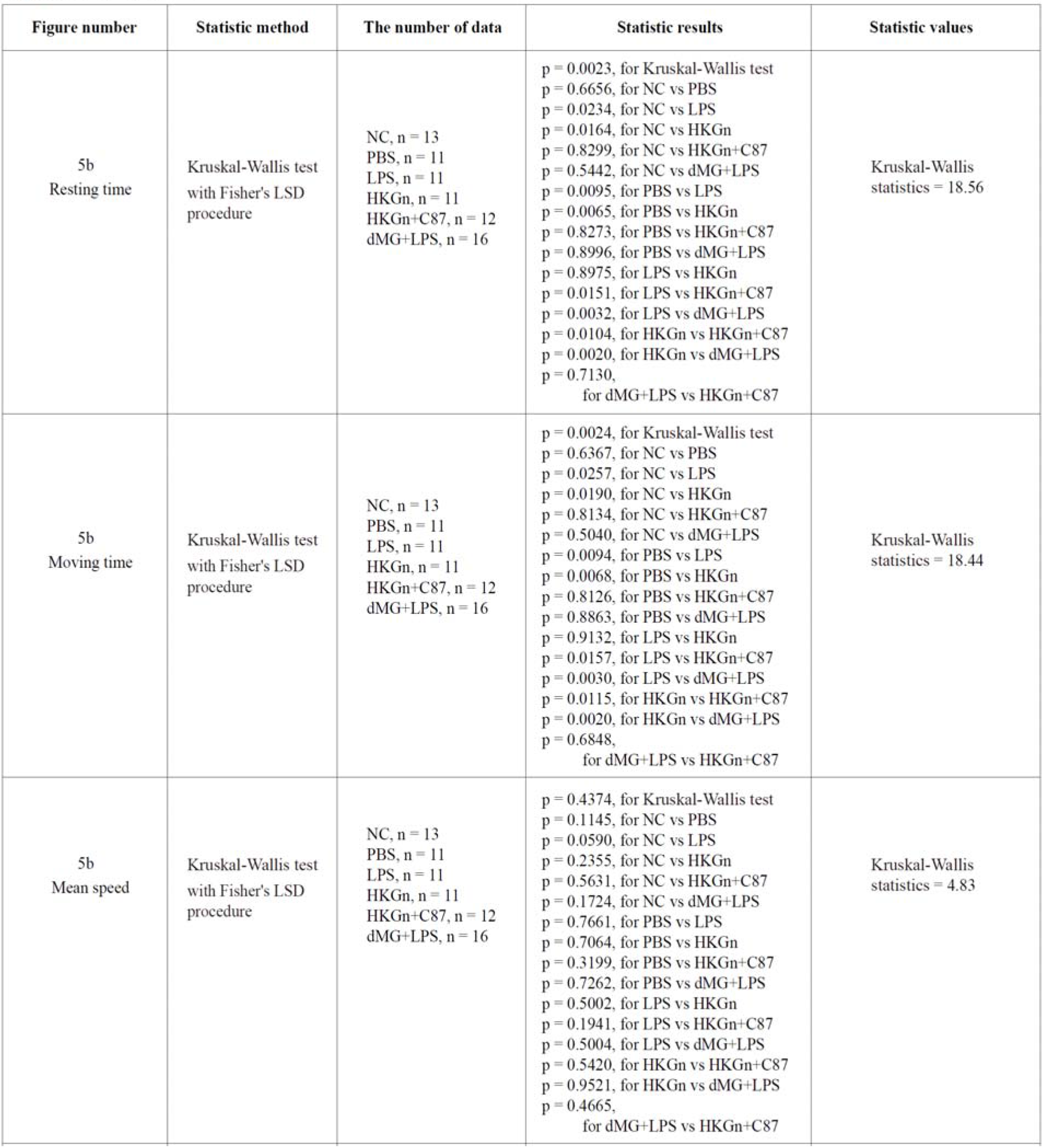

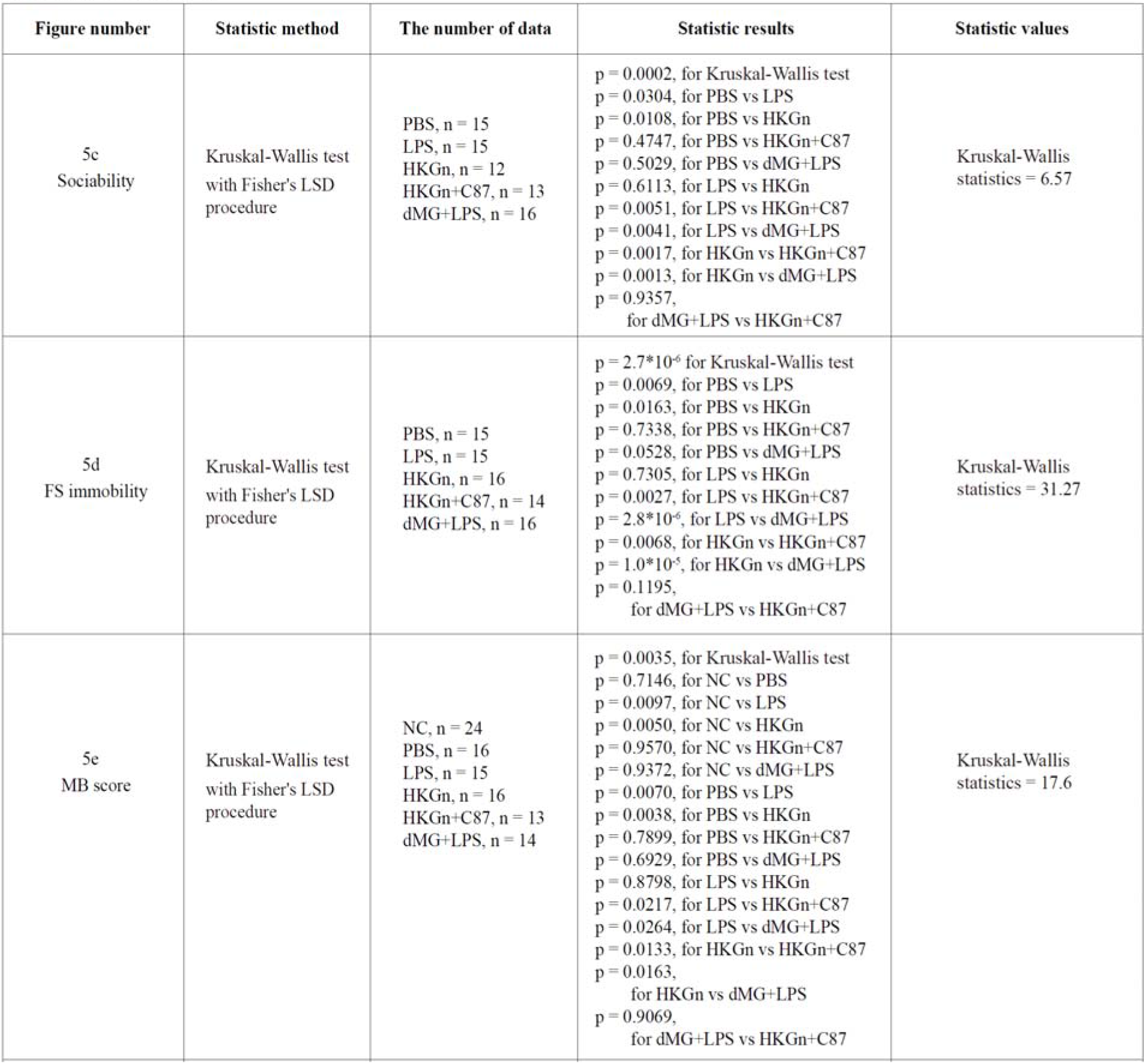

**Supplementary Figure 1.**
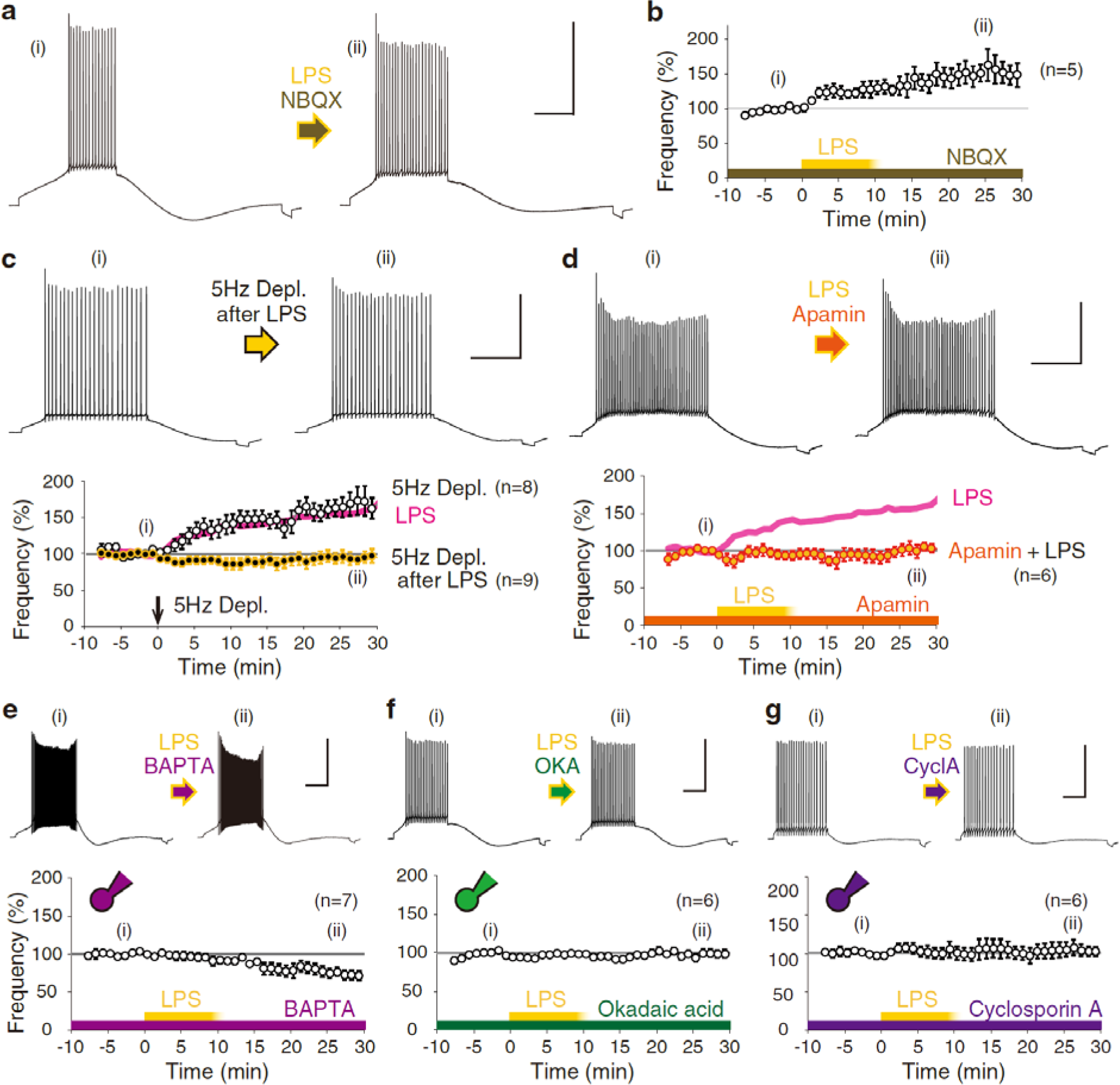
Signaling involved in LPS-induced intrinsic plasticity in Purkinje neurons. **a,** Representative action potential firing of Purkinje neurons before and after LPS (lipopolysaccharide, 10-12 μg/mL) treatment in the presence of an AMPA (α-amino-3-hydroxy-5- methyl-4-isoxazolepropionic acid) receptor blocker, NBQX (20 μM) (*p < 0.03, Mann-Whitney *U*-test). **b,** Time course of the change in firing frequency normalized between −5 to −1 min. LPS application began at 0 min and continued for 10 min. **c,** Occlusion of the increase in firing plasticity in Purkinje neurons pre-exposed to LPS. Time courses of the normalized frequency of neurons following 5-Hz depolarization and that after 15-minutes LPS exposure. Conditioning was applied at 0 min, as indicated by the arrowhead. LPS-only data are superimposed for comparison (magenta line) (5Hz Depl.+LPS, p > 0.8; 5Hz Depl., *p < 0.005). **d,** Suppression of LPS-induced increases in firing frequency under treatment with the SK channel blocker, apamin (10 nM). A time course of the normalized frequency of neurons treated with LPS in the presence of apamin is shown. LPS application began at 0 min. **e-g,** Intra-neuronal administration of BAPTA (20 mM, e), okadaic acid (OKA, 150 nM, f) and cyclosporin A (CycloA, 100 μM, g) inhibited increases in firing frequency by LPS exposure (BAPTA, ^†^p < 0.01 for reduction; OKA, p > 0.4; CycloA, p > 0.5). Results suggest that LPS-induced increases in firing frequency were dependent on Ca^2+^, PP1, PP2A, and PP2B. Drug application began at 0 min. Scale: 40 mV and 200 ms. LPS, lipopolysaccharide.

**Supplementary Figure 2.**
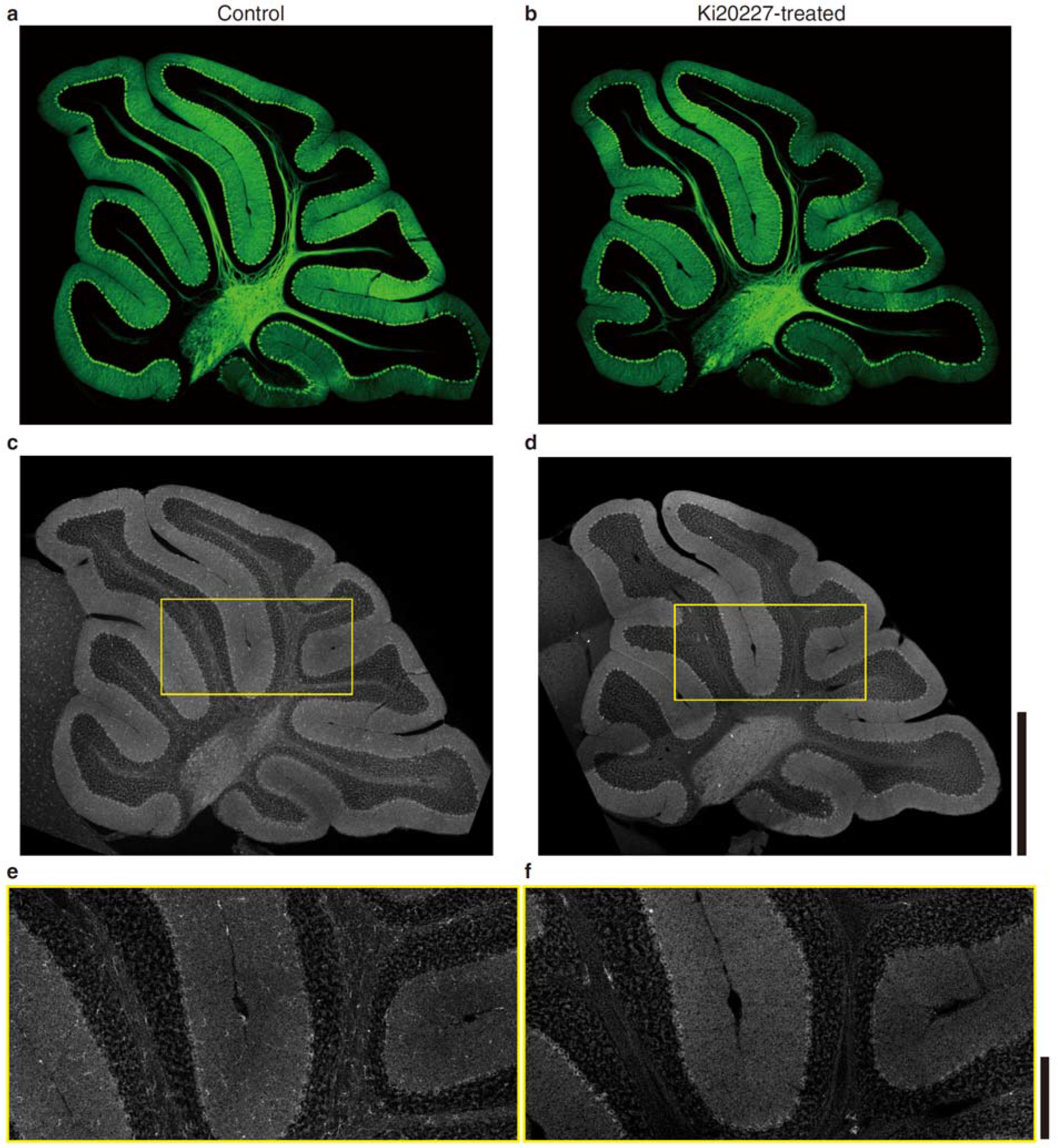
Microglia depletion in the cerebellum of Ki20227-treated mice. Whole cerebellar sagittal images of control (a and c) and Ki20227-administered (b and d) mice by calbindin (green) and Iba1 (white) co-immunostaining. Expansion images of yellow squares are shown in bottom (e and f, respectively). Note that the Iba1 positive microglia are not present in the Ki20227-administered cerebellum. Bar scales are 1 mm (c, d) and 200 μm (e, f).

**Supplementary Figure 3.**
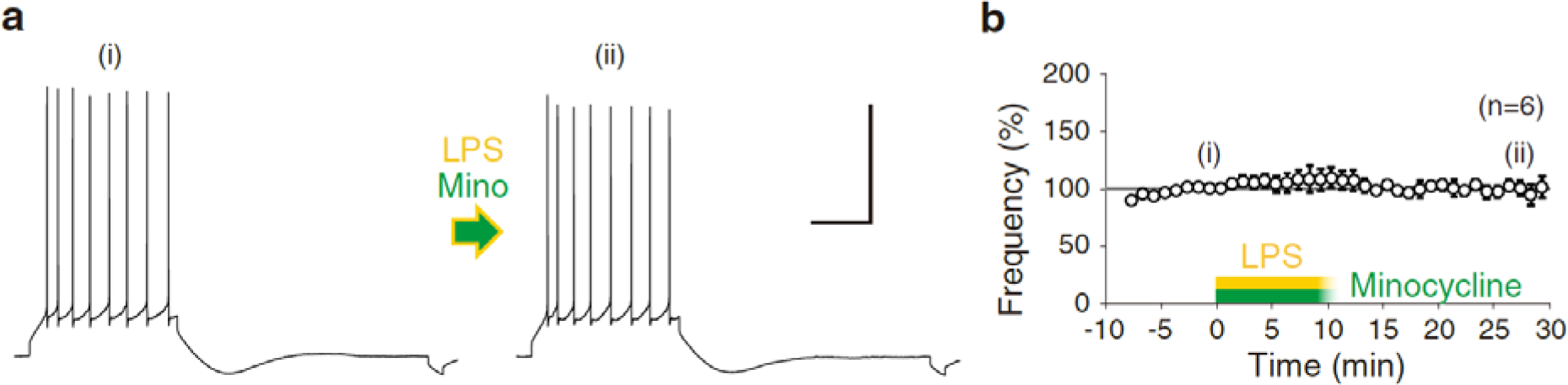
Abolishment of LPS-induced intrinsic plasticity under suppression of microglia. **a,** Action potential firing of Purkinje neurons before and after exposure to LPS (12 μg/mL) with minocycline-hydrochloride (50 nM). **b,** Time course of the normalized firing frequency (Minocycline+LPS, p > 0.7, Mann-Whitney *U*-test). Drug application began at 0 min. Scale: 40 mV and 200 ms.

**Supplementary Figure 4.**
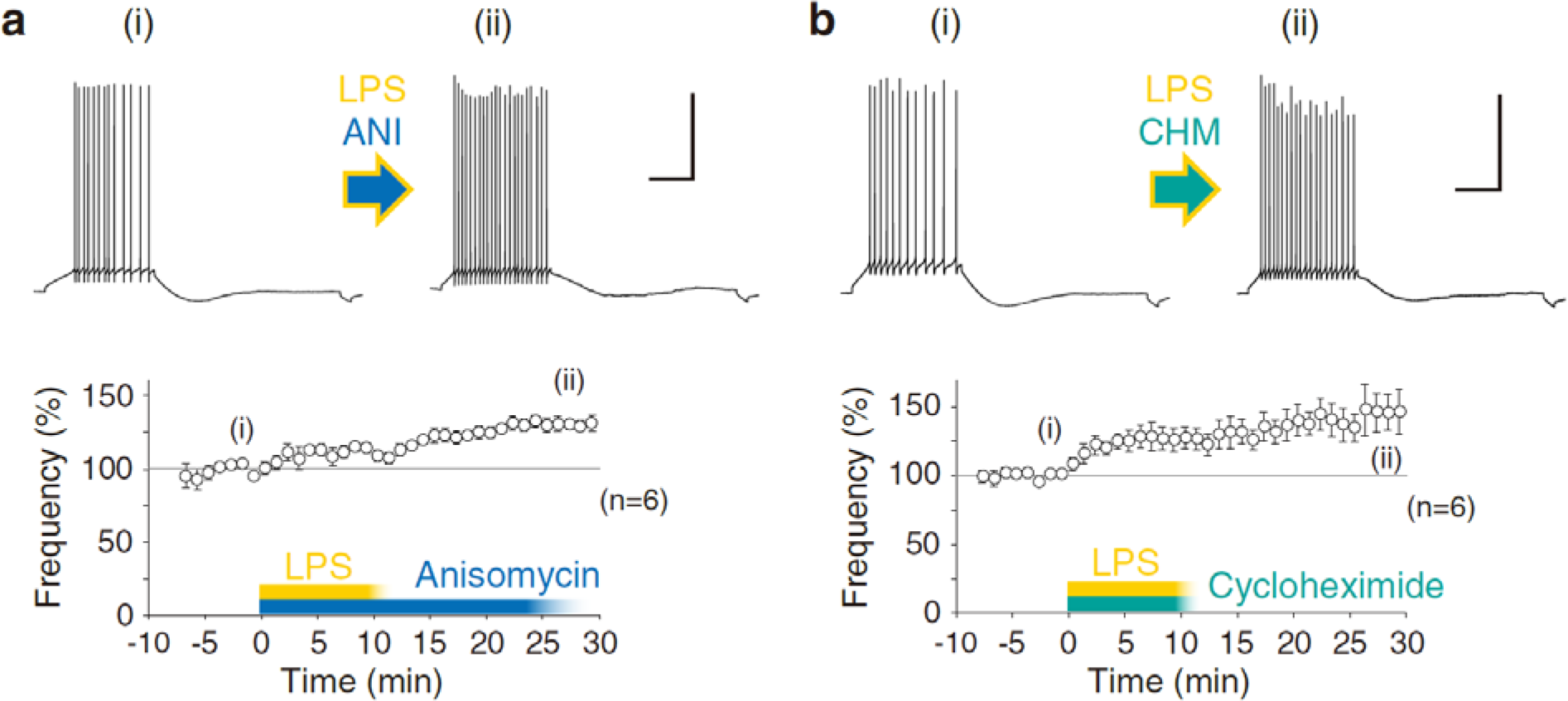
Protein-synthesis independence of LPS-induced excitability plasticity. **a,** Action potential firing of Purkinje neurons following LPS exposure under treatment with anisomycin (30 μM) (*p < 0.03, Mann-Whitney *U*-test). **b,** Action potential firing following LPS exposure under treatment with cycloheximide (30 μM) (*p < 0.03). While both data suggested that the firing increase by exposure to LPS was induced under suppression of protein translation, a minor, and not significant, reduction of the extent of increase in firing frequency was notified compared to LPS-conditioned experiments as shown in Fig.1b. Scale: 40 mV and 200 ms. LPS-application began at 0 min and continued for 10 min.

**Supplementary Figure 5.**
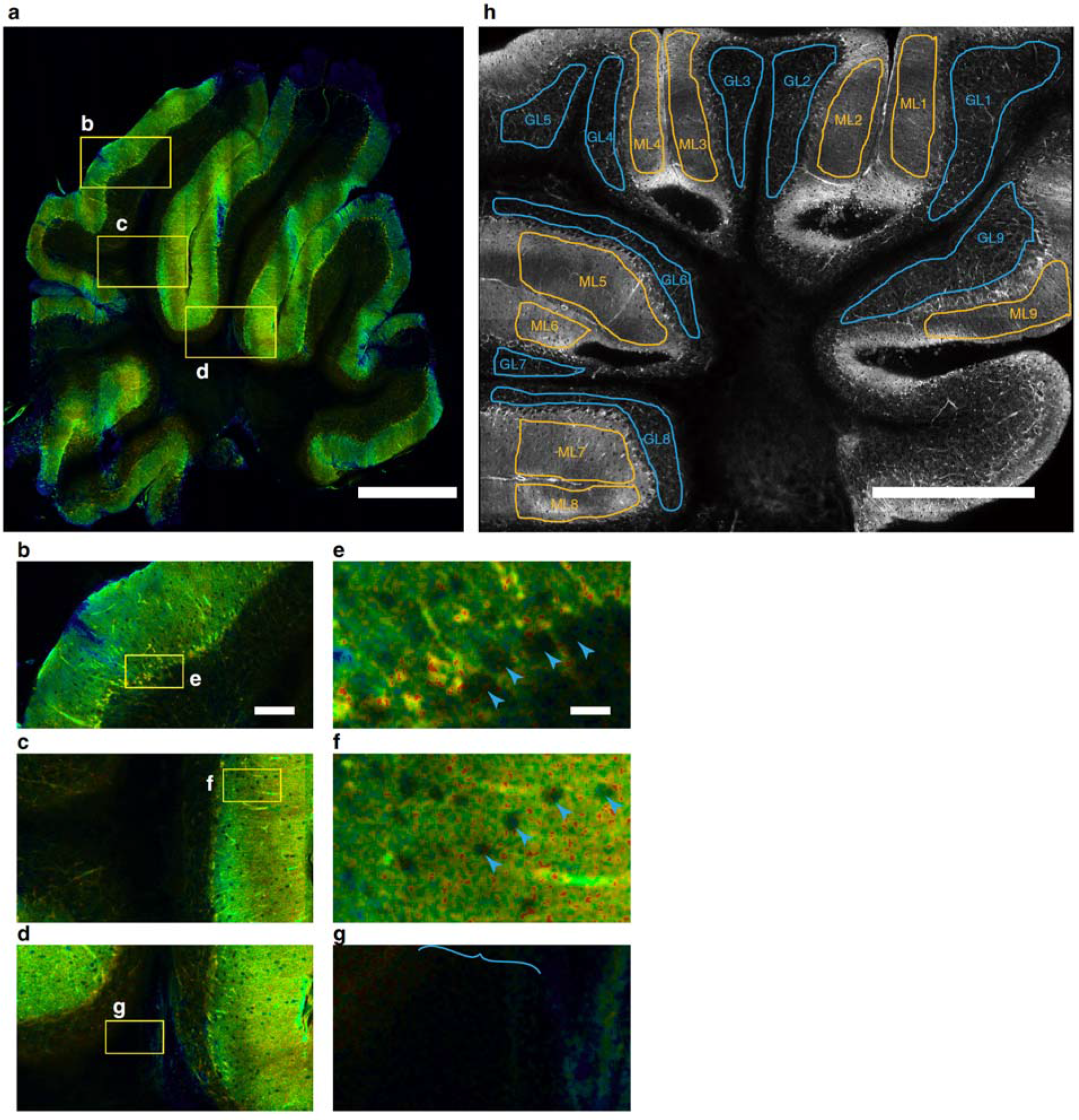
FRET image of a whole cerebellar sagittal section. **a,** A merged tile image of GO-ATeam2 cerebellum is shown as a sagittal section. Please note the vertical stripe of discontinuity is due to the slice anchor. **b-d,** Expanded images in the yellow flamed regions in (a). **e-g,** Expanded images in the yellow flamed regions in (b-d), respectively. Arrowheads indicate the cell body of Purkinje neurons (e), or interneurons in the ML (f). The curly bracket indicates the location of a bundle in the white matter (g). **h,** ROIs of ATP concentration changes were selected as in the image. Numbering of ML-X and GL-X indicates in the molecular layer and in the white matter, respectively. Bar scale: 1 mm (a, h), 200 μm (b), and 40 μm (e). FRET, fluorescence/Förster resonance energy transfer.

**Supplementary Figure 6.**
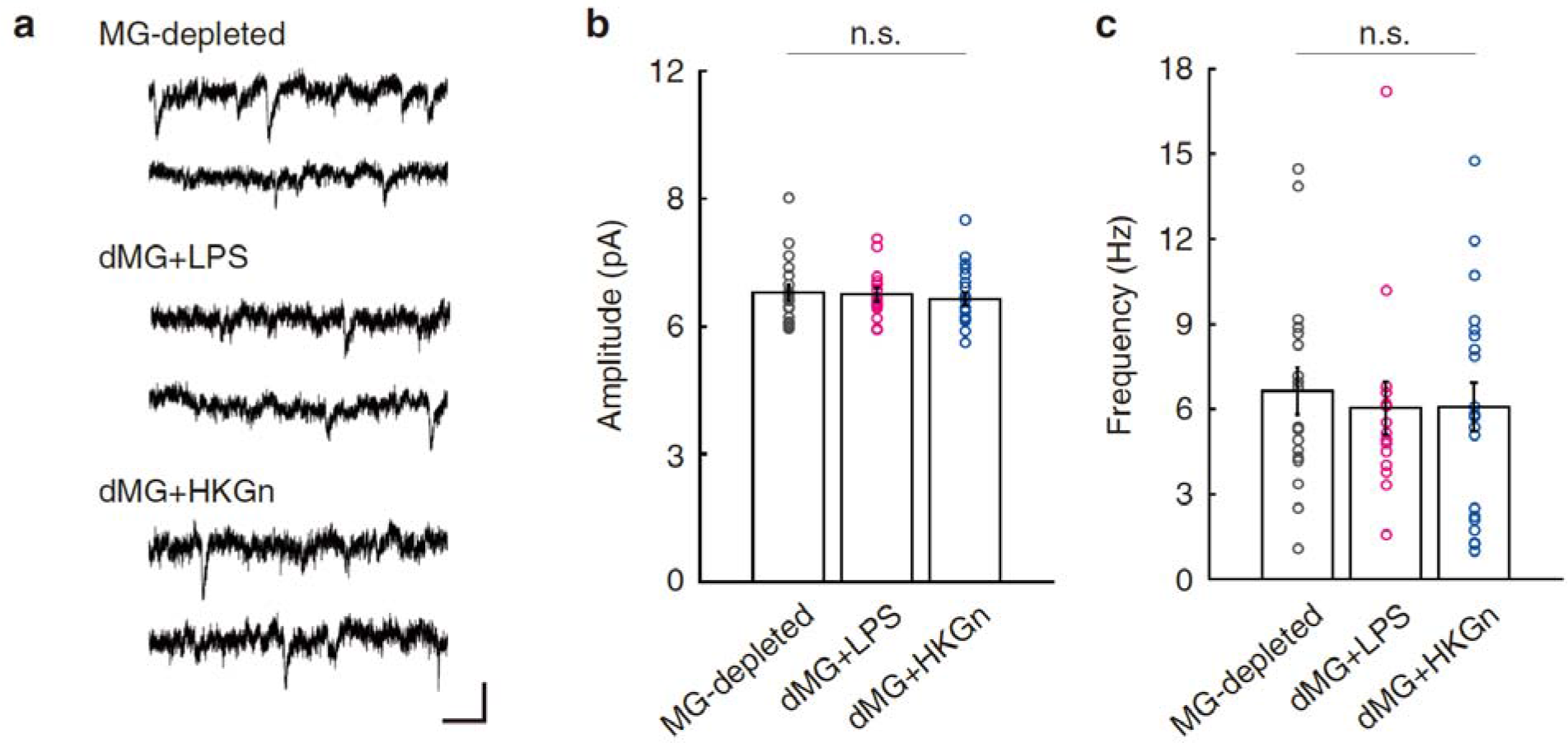
Excitatory synaptic transmission in Purkinje cells in microglia depleted cerebella. **a,** Representative spontaneous excitatory postsynaptic currents (sEPSCs) traces of microglia (MG)-depleted mice. Scale: 10 pA, 100 ms. Bar graphs of amplitude (b) and frequency (c) of sEPSC in different experiments (mean ± SEM) are shown. Total numbers of 18, 15 and 21 cells were recorded in the MG-depleted (dMG) control, dMG LPS- and HKGn-exposure, respectively. Amplitude: MG-depleted, 6.8 ± 0.2 pA; dMG+LPS, 6.8 ± 0.2 pA; dMG+HKGn, 6.6 ± 0.2 pA. Frequency: MG-depleted, 6.7 ± 0.8 Hz; dMG+LPS, 6.0 ± 0.9 Hz; dMG+HKGn, 6.1 ± 0.8 Hz. No significant differences (n.s.) were observed among all pairs (p > 0.3, in Mann-Whitney *U*-test against MG-depleted control), as well as those in rise time, half-width, and decay time (*data not shown*).

**Supplementary Figure 7.**
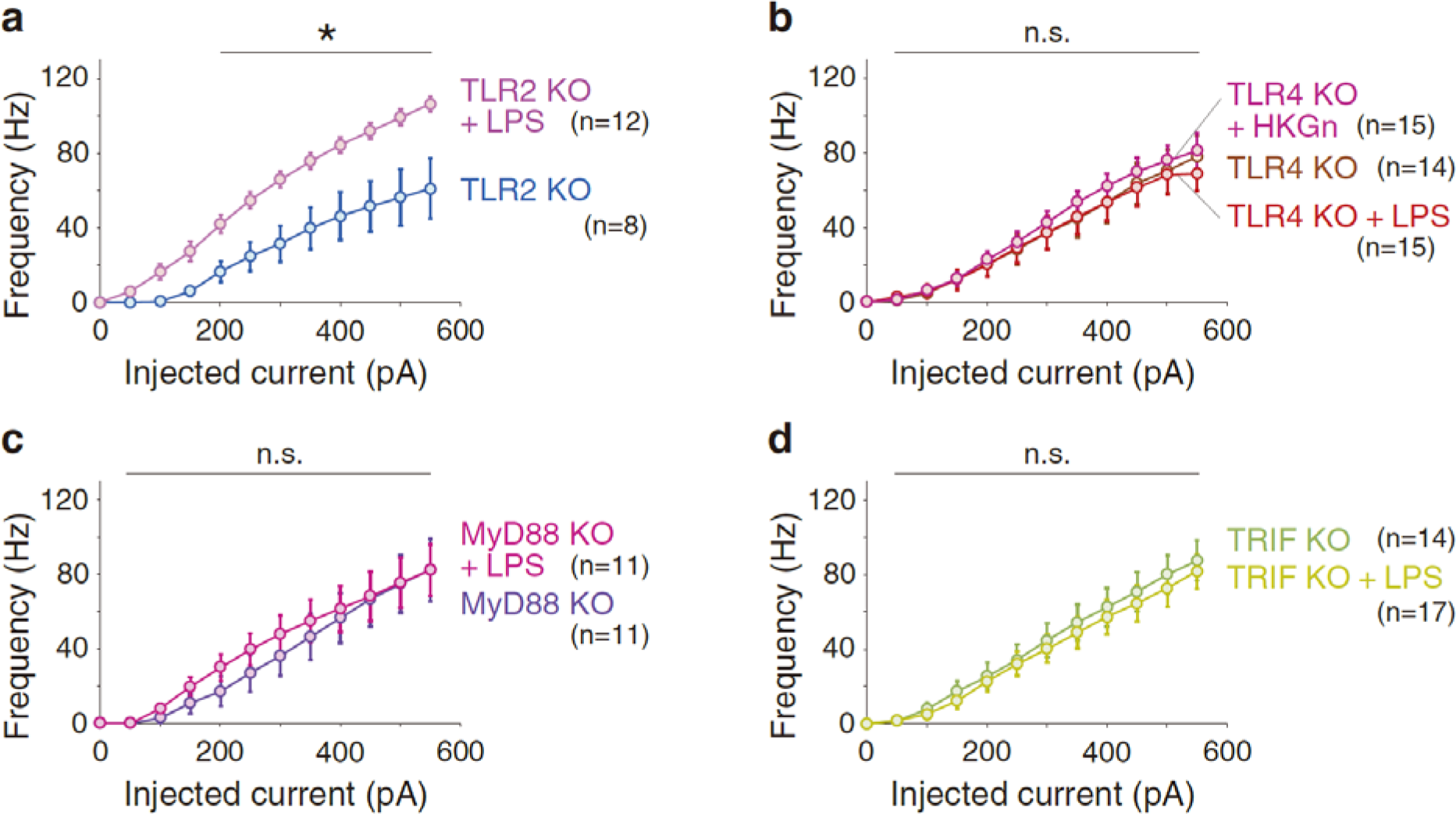
Firing frequency of TLRs, MyD88 and TRIF KO Purkinje cells. Firing frequency of TLR2 KO (a), TLR4 KO (b), MyD88 KO (c) and TRIF KO (d) neurons in response to different depolarization pulses. No significant difference was observed between LPS- or HKGn-exposed and non-exposed neurons in either transgenic mouse line, except for in TLR2 KO (a). Asterisk indicates p < 0.05 in the Mann-Whitney *U*-test. MyD88 KO, Myeloid differentiation primary response gene 88 knockout mice; TLR2/4 KO, Toll-like receptor 2/4 knockout mice; TRIF KO, TIR-domain-containing adapter-inducing interferon-β knockout mice.

**Supplementary Figure 8.**
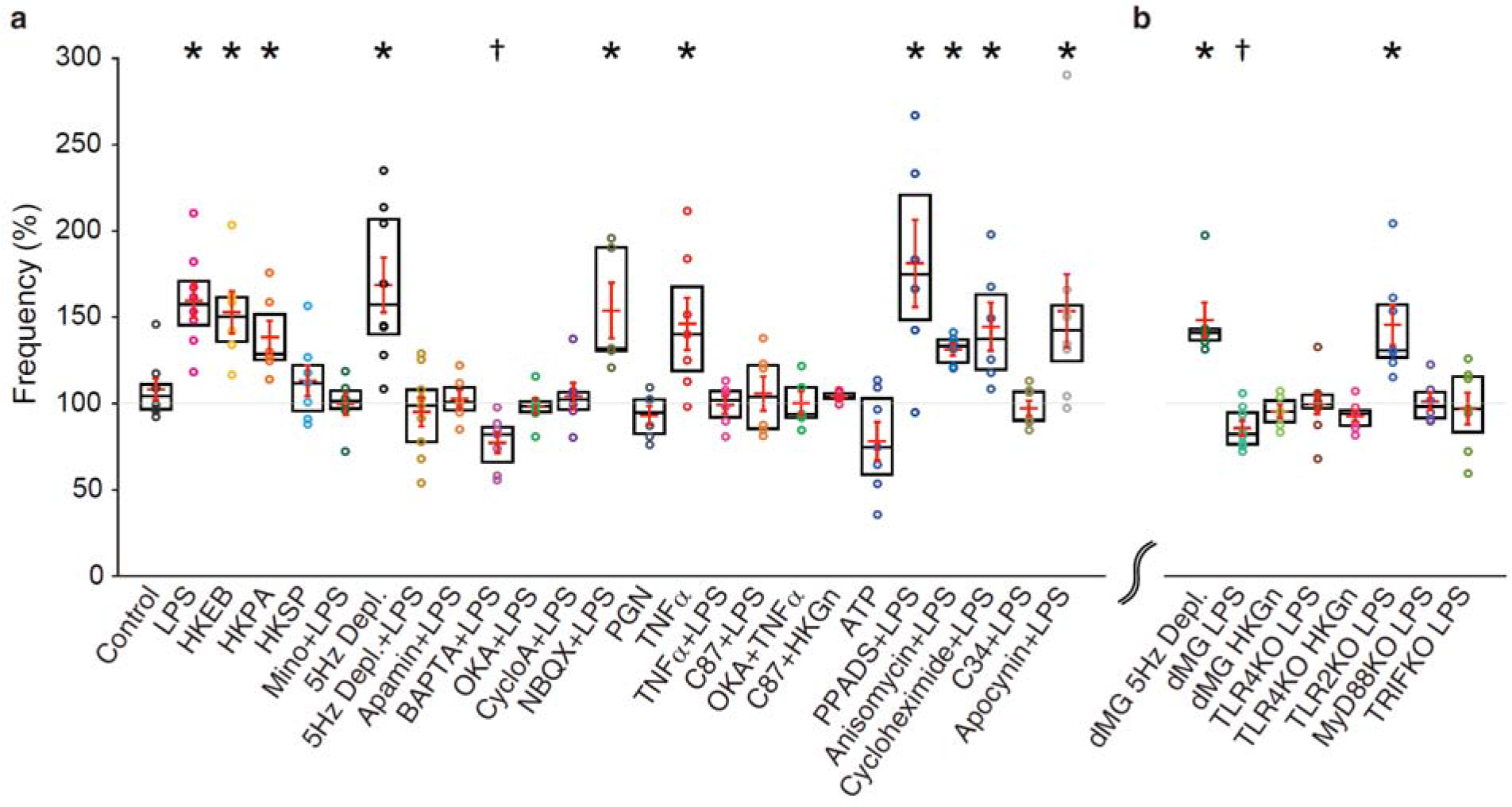
Summary of the extent of excitability changes. Firing frequency changes after the application of various reagents or conditioning for intrinsic plasticity in rat (a) and mouse neurons (b). All data from long-term recordings in this study were summarized as the mean ± SEM (red mark) with a box plot. Asterisks indicate significant increases (p < 0.05, in the Mann Whitney *U*-test). Daggers indicate significant decreases (p < 0.05).

**Supplementary Figure 9.**
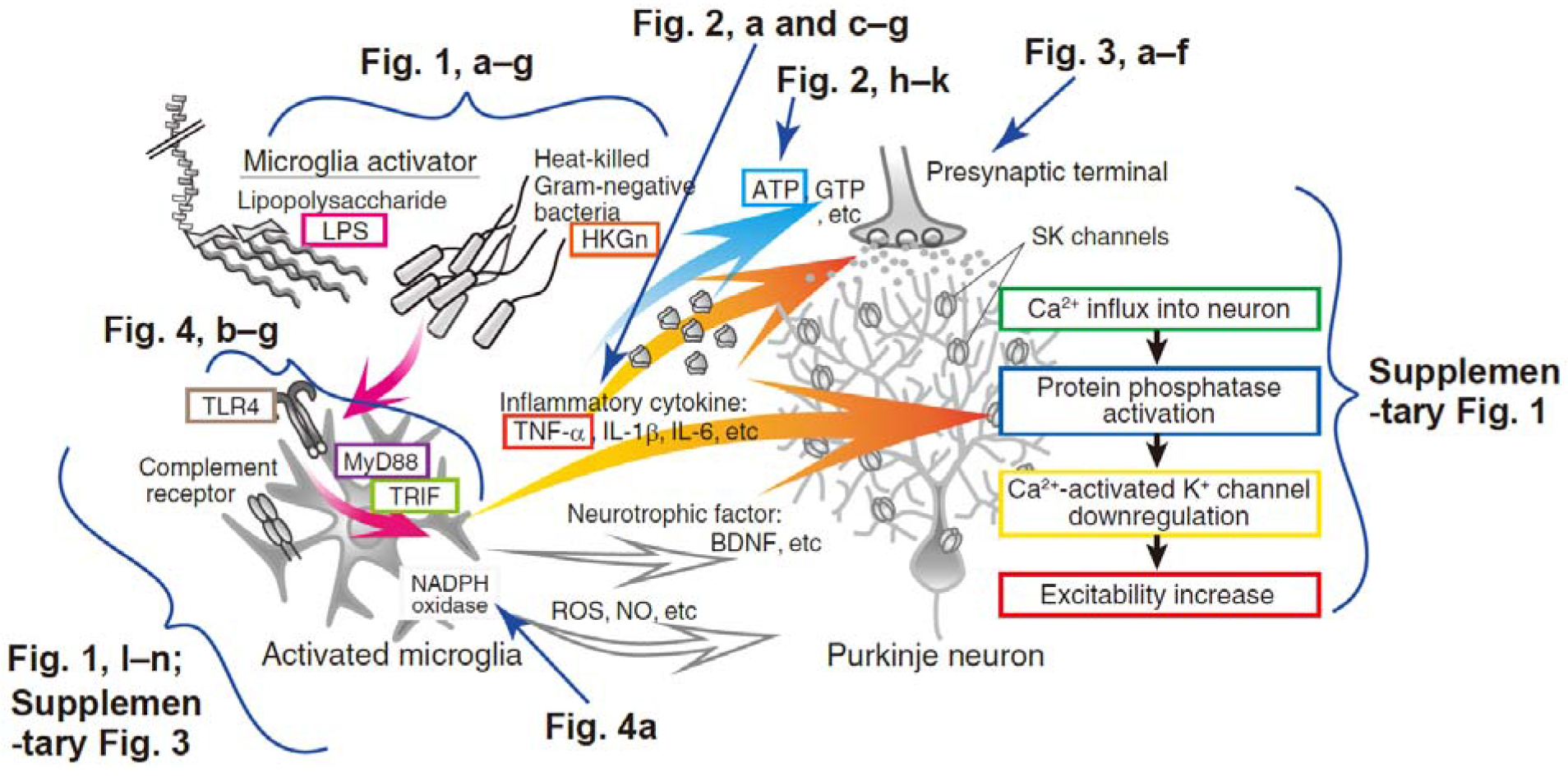
Experimental diagram of the endotoxin-induced excitability plasticity. Corresponding figure numbers are tagged or arrowed to the signaling cascade.

**Supplementary Figure 10.**
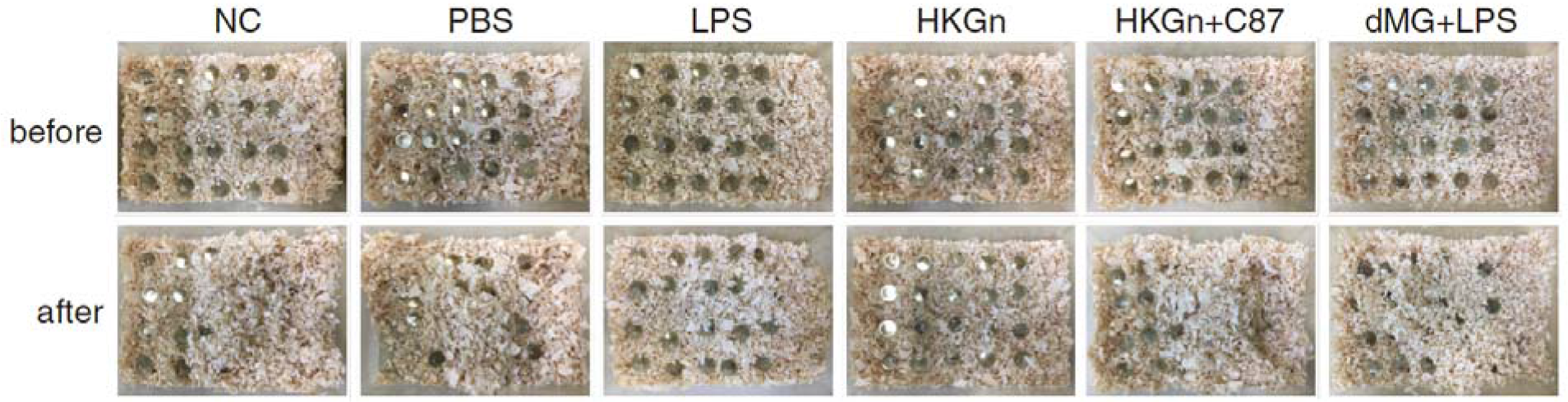
Marble burying of rats injected microglia activators and those suppressed inflammation. Pictures of marble burying behavior before and after 20-minutes monitoring of drug-infused rats. PBS, LPS, heat-killed Gram-negative bacteria mixture (HKGn: HKEB+HKPA), HKGn+C87, or nothing (NC) was injected into the cerebellar vermis. In other experiments, LPS was injected into the cerebellum of microglia-depleted rats (dMG+LPS). Note that rats with LPS- and HKGn-injection hid less marbles compared to NC, PBS, HKGn+C87 and dMG+LPS.

**Supplementary Figure 11.**
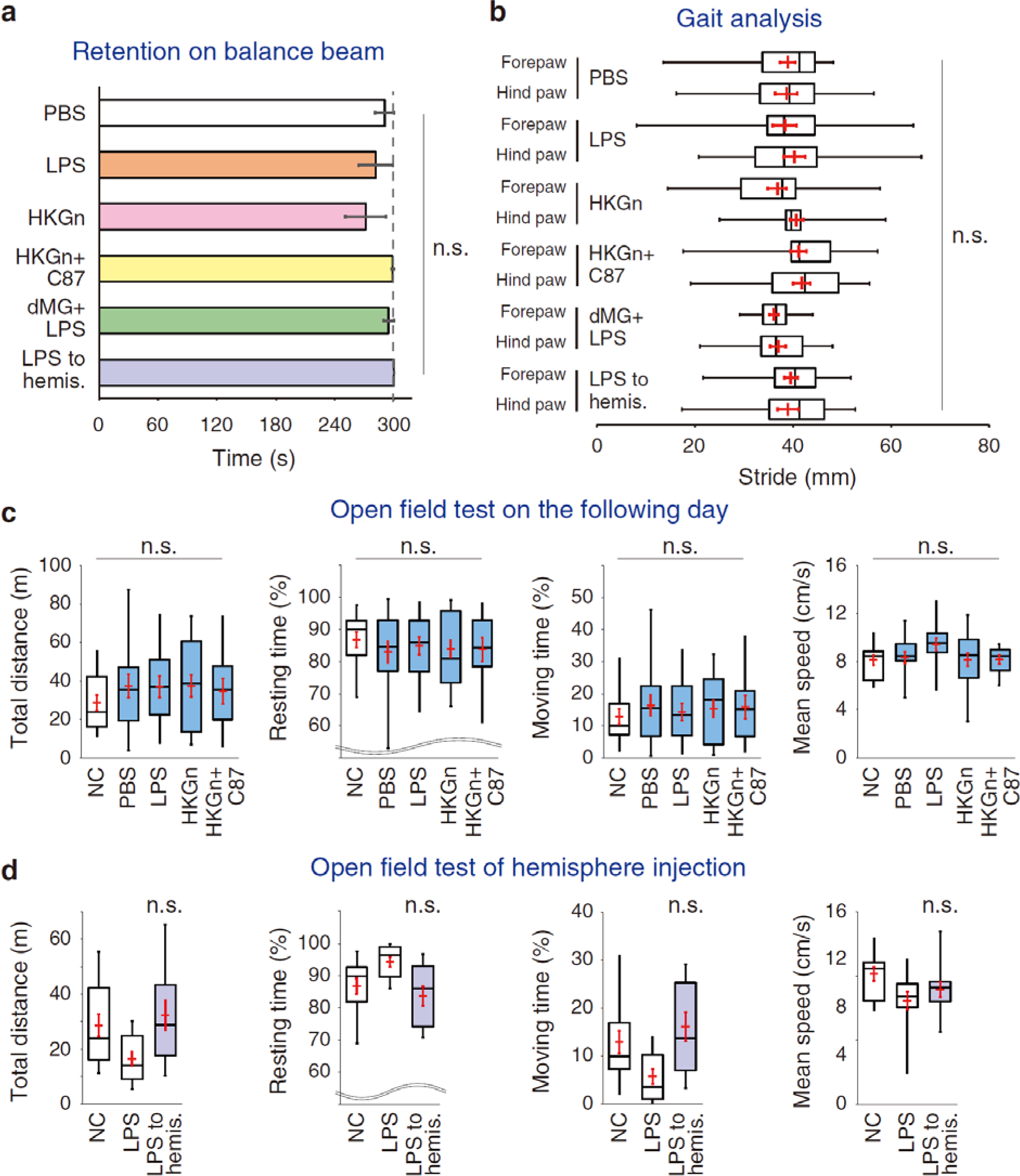
Other behavior paradigms of reagent-injected animals. **a,** Retention time on balance beam of drug-infused rats in total 5-minute measurement. PBS, LPS, heat-killed Gram-negative bacteria mixture (HKGn), and HKGn+C87 was injected into the cerebellar vermis. In some experiments, LPS was injected into the cerebellum of microglia-depleted rats (dMG+LPS). Alternatively, LPS was injected into the cerebellar hemispheres (LPS to hemis.). **b,** Stride length during walking in reagent-infused rats. **c,** Exploration behavior in the open field arena one day after drug-infusion. Box plots of the total distance, resting time, moving time, and mean speed of PBS-, LPS-, HKGn- and HKGn+C87-infused rats on the following day are shown as light-blue colored boxes. **d,** Open field test of LPS to hemispheres-animals. White boxes in (c) and (d) are NC and LPS data of Fig. 5b for comparison. No significant differences were found in all parameters.

**Supplementary Figure 12.**
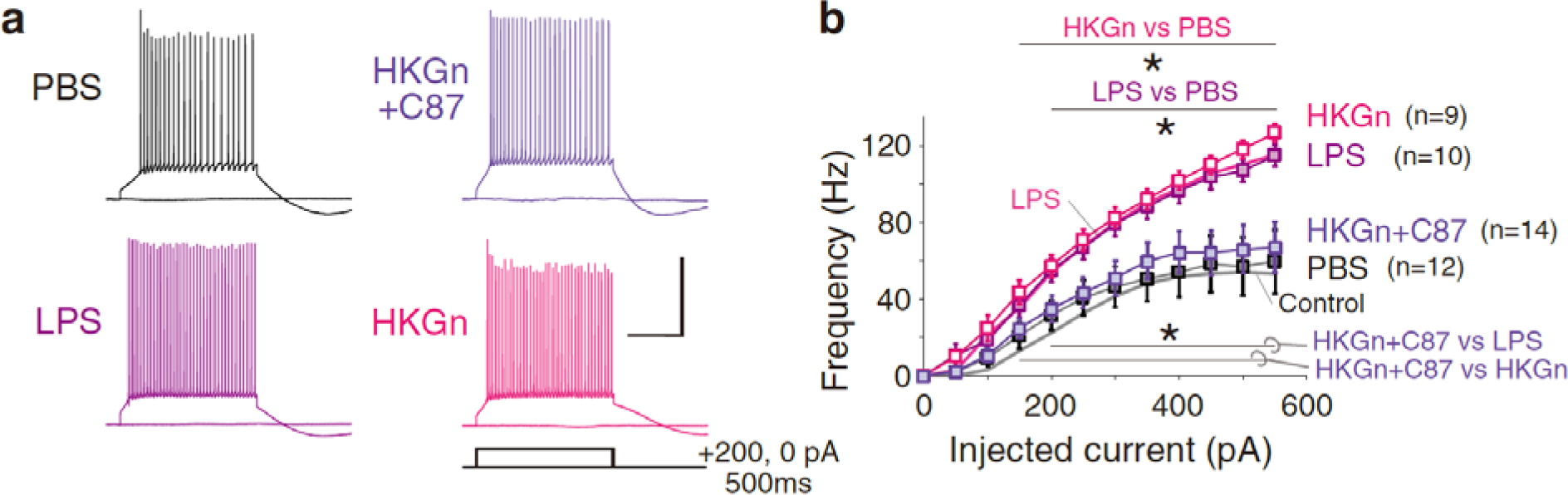
Firing frequency of Purkinje cells obtained from drug-treated rats. Representative traces (a) and the firing frequency (b) of neurons under slice preparation obtained from the cerebellar vermis of rats injected with control saline, LPS, heat-killed Gram-negative bacteria (HKGn), or HKGn+C87 in response to different depolarization pulses. Mean firing frequency of LPS-exposed and control neurons from Fig. 1b are merged for comparison (b). Neurons from LPS- and HKGn-injected cerebella exhibited a significant increase in firing frequency relative to that observed for control saline-injected and HKGn+C87 co-injected neurons (*p<0.05), suggesting that the TNF-α inhibitor suppressed the increase in Purkinje cell excitability *in vivo.* Scale: 40 mV and 200 ms. TNF-α, tumor necrosis factor alpha

**Supplementary Figure 13.**
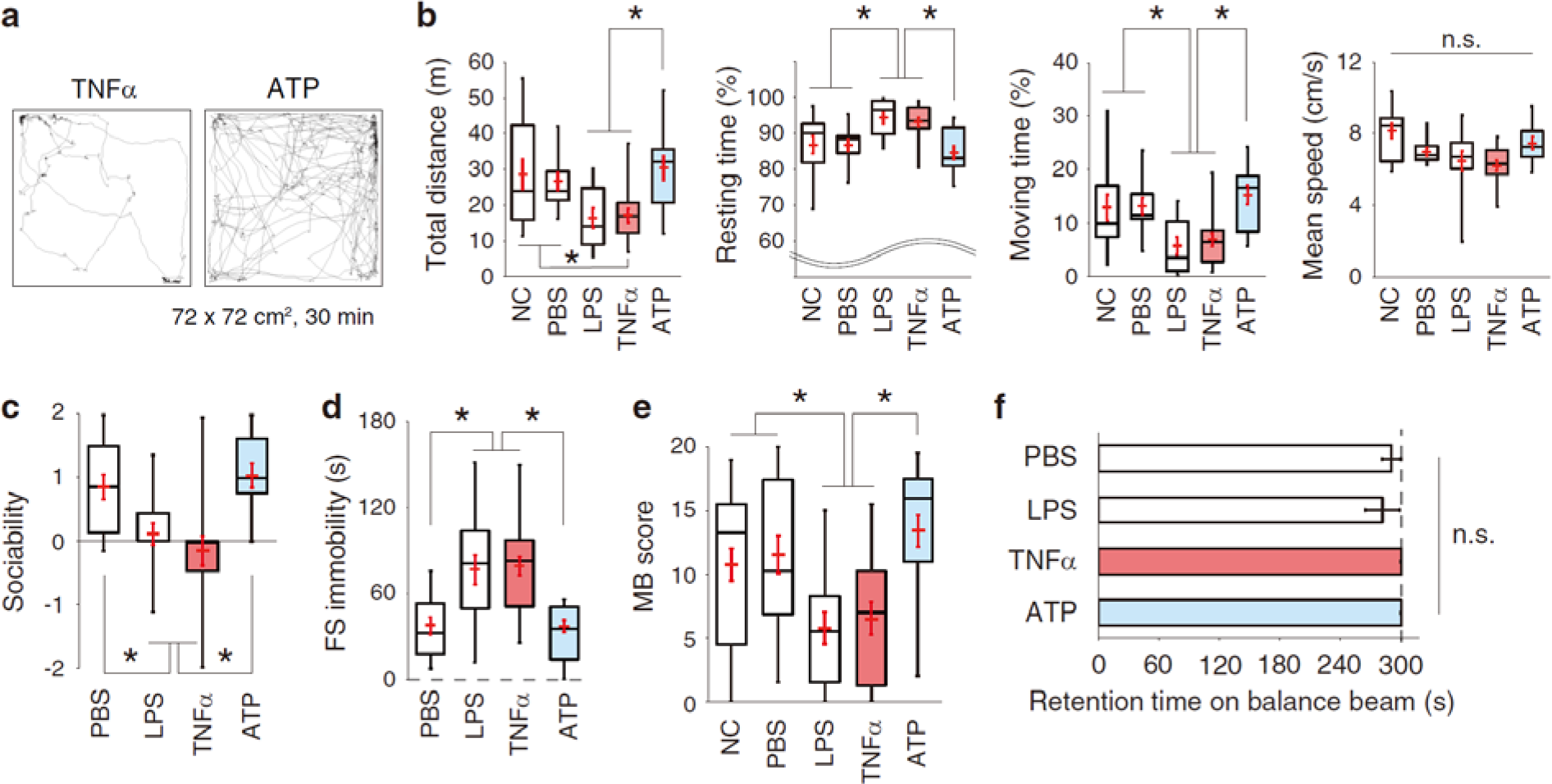
Behavior tests in ATP and TNF-α-infused animals. **a,** Representative trajectories of reagent-infused rats in the open field arena. TNF-α (20 μg/mL) or ATP (20 mM) was injected into the anterior cerebellar vermis. **b,** Box plots of the total distance, resting time, moving time, and mean speed. Injection of TNF-α, but not ATP, significantly reduced the animals’ exploratory behavior, sociability, forced motivation and repetitive behaviors. **c,** Social interaction. **d,** Forced swim test. **e,** Marble burying. **f,** Posture retention test on balance beam. White boxes in (b-f) are NC, PBS and LPS data of Fig. 5b-e and Supplementary Fig. 12b for comparison. Overlapping red marks represent mean ± SEM. *p < 0.05, *multiple comparison*.

**Supplementary Figure 14.**
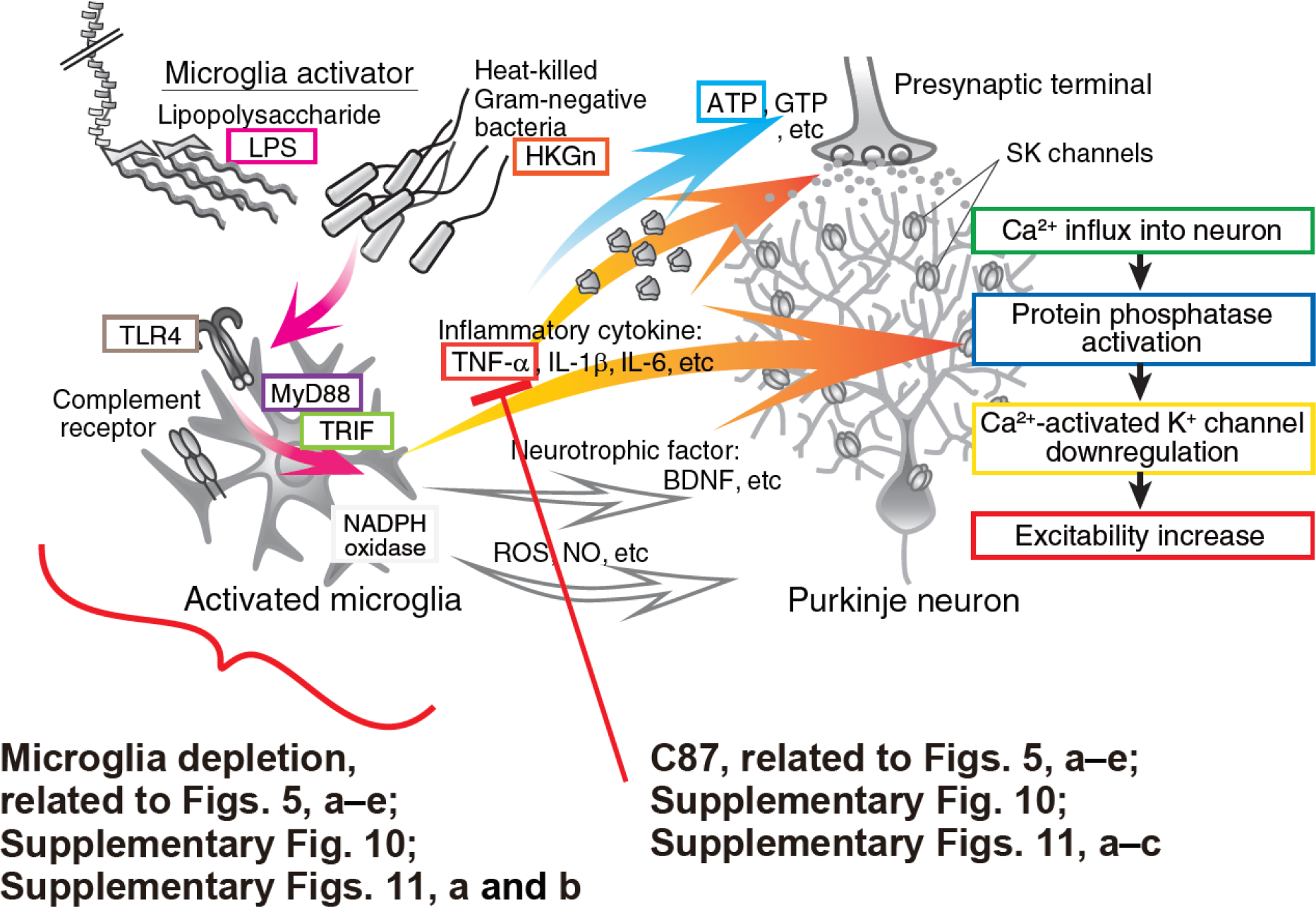
Potential targets for the rescue of behavior abnormality by cerebellar inflammation. The depletion of microglia and inhibition of TNF-α by C87 used in this study are shown.

**Supplementary Figure 15.**
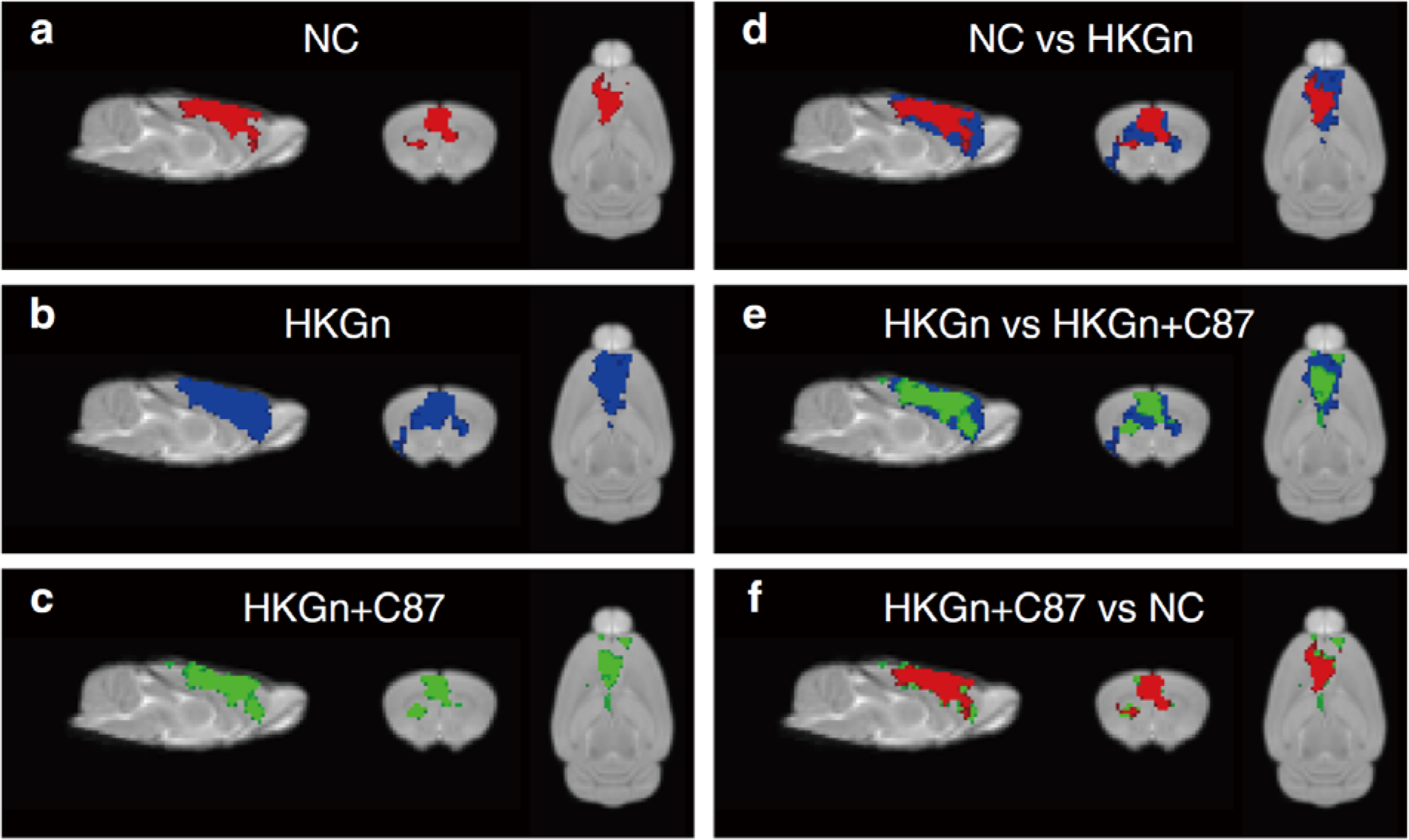
Components of frontal cortical area from group ICA of drug treated rats. Group-based ICA spatial maps are shown in the same component of NC (red), HKGn (blue) and HKGn+C87 (green) (a-c, respectively). Merged maps of NC vs HKGn, HKGn vs HKGn+C87 and HKGn+C87 vs NC are shown in d-f, respectively. Images represent spatial color coded 1-p maps of the component, overlaid onto anatomical T2W image. Images were thresholded by p<0.01. Of note, animals HKGn-infused to the cerebellar anterior vermis express larger area of activation in the medial prefrontal cortex (including infralimbic and prelimbic cortices), cingulate cortex and primary motor area. And, the enlargement of frontal areas was recovered by C87 administration in the cerebellum. Sagittal, coronal, and transverse planes of brain images are shown (from left to right). ICA, independent component analysis; NC, non-conditioned; HKGn, heat-killed Gram-negative bacteria mixture.

## Supplementary information

### Video 1

Representative movies of open field test of control, inflammation-induced, and rescued animals. Movies fast-forwarded to ×32 are arranged. From the up-left to up-right: No-conditioned, LPS- injected rats, and a Ki20227-administered rat with LPS injection. From the bottom-left to bottom-right: PBS-injected, HKGn-injected, and HKGn+C87-injected rats. (Related to Fig. 5a,b).

